# Biogenesis of specialized lysosomes in differentiated keratinocytes relies on close apposition with the Golgi apparatus

**DOI:** 10.1101/2022.12.22.521712

**Authors:** Sarmistha Mahanty, Ptissam Bergam, Vivek Belapurkar, Litralson Eluvathingal, Nikita Gupta, Bruno Goud, Deepak Nair, Graca Raposo, Subba Rao Gangi Setty

**Author notes:** **Corresponding authors:** Sarmistha Mahanty - *Subba Rao Gangi Setty. Foot notes: Authors contributed equally to this work Lead contact (Graca Raposo, Subba Rao Gangi Setty).

## Abstract

Intracellular organelles support cellular physiology in diverse conditions. In the skin, epidermal keratinocytes undergo differentiation with gradual changes in cellular physiology, accompanying remodeling in lysosomes and the Golgi apparatus. However, the functional significance and the molecular link between lysosome and Golgi remodeling were unknown. Here, we show that in differentiated keratinocytes, the Golgi apparatus redistributes as ministacks leading to a significant increase in total protein secretion. The Golgi ministacks establish contact with lysosomes facilitated by Golgi tethering protein GRASP65, the depletion of which is associated with the loss of Golgi-lysosome contact and malformation of lysosomes defined by their aberrant morphology, size, and function. Strikingly, these lysosomes receive secretory Golgi cargoes, contribute to the protein secretion from the Golgi, and are critically maintained by the secretory function of the Golgi apparatus. We uncovered a novel mechanism of lysosome specialization through unique Golgi-lysosome contact that likely supports high secretion from differentiated keratinocytes.

**Key points:**

- Calcium induced differentiation of human keratinocytes accompanies the dispersal of functional Golgi stacks and lysosomes.
- Dispersed Golgi stacks establish contact/physical apposition with the lysosomes.
- Golgi tether GRASP65 surrounds keratinocyte lysosomes and facilitates Golgi-lysosome apposition.
- GRASP65 depletion abolishes Golgi-lysosome apposition and accumulates morphologically altered non-degradative lysosomes.
- Lysosomes of differentiated keratinocytes receive secretory Golgi cargo and contribute to the protein secretion from the Golgi.
- Specialized lysosomes are maintained by the secretory function of the Golgi apparatus.

## Introduction

Eukaryotic cells are compartmentalized into membrane-bound autonomous intracellular organelles. However, organelle function and cellular homeostasis often require communication between organelles via physical apposition or contact (between 20 - 40 nm) that facilitates the exchange of materials such as signaling molecules, Ca^2+^ and lipids ^1^. For instance, lysosomes establish contacts with mitochondria and endoplasmic reticulum (ER) that regulate cellular Ca^2+^ homeostasis and Ca^2+^ transport to the lysosomes required for their proper functioning and acidification, respectively ^2,3^. Lysosome-peroxisome contacts regulate lysosome mediated lipid homeostasis ^4,5^. The biogenesis of lysosomes also depends on late endosomes and the Golgi apparatus. Lysosomes receive luminal content from late endosomes, and newly synthesized lysosomal hydrolases and membrane proteins are transported from the Golgi ^6,7^. Several secretory organelles including the Lysosome-related organelles (LROs) such as Weibel-Palade bodies (WPBs) and neuronal secretory granules, originate from the Golgi and mature through subsequent endosomal input ^8^. However, whether the Golgi directly contributes to the maturation of classical lysosomes and whether as yet undescribed physical contacts between lysosomes/late endocytic organelles with the Golgi play a role in lysosome maturation is not understood.

The Golgi apparatus in mammalian cells is a perinuclearly localized compact structure consisting of multiple stacked cisternae that are arranged from *cis* to *trans* as a ribbon. The ribbon structure favors a complex, stepwise glycoprotein processing ^9, 10^. However, the Golgi apparatus disassembles and disperses under some physiological conditions to accommodate higher secretion rate or during mitosis to facilitate Golgi segregation between daughter cells^11^. For instance, dispersed Golgi stacks are observed in prolactin-secreting cells of female rats following induction by the sucking stimulus (Rambourg et al., 1993). In neurons and in cells of the platypus Brunner’s gland, Golgi outposts are formed to facilitate localized protein transport ^12–15^. The dynamic morphology and function of the Golgi are regulated by Golgi-associated structural and functional proteins, such as the Golgi reassembly stacking proteins GRASP55 and GRASP65 and the small GTPase, ADP-ribosylation factor1 (ARF1) ^16,17^. GRASP55 and GRASP65 play essential and complementary functions in maintaining Golgi ribbon morphology by linking the Golgi stacks through their oligomerization regulated by cyclin-dependent kinases (CDKs) and polo-like kinases ^9,18,19^. The small GTPase ARF1 regulates intra-Golgi and Golgi-ER transport by the recruitment of COPI (coat protein I complex) to form COPI vesicles ^16,20^. An increase in the secretory function requires an increase in the generation of COPI vesicles, which can be positively regulated by the dispersion of Golgi stacks, as shown previously in an *in vitro* budding assay ^21^.

Recent studies have established a number of molecular links between the functions of the Golgi complex and lysosomes. For example, the master growth-regulating kinase, mTORC1, predominantly localizes to lysosomes where it can respond to changes in cellular metabolism, but a cohort of mTORC1 also localizes to Golgi membranes and is activated through the Golgi localized small GTPase Rab1 and the Golgi membrane tether GOLPH3 ^22–24^. Furthermore, the perinuclear localization of lysosomes during starvation is influenced by the interaction between the Golgi-localized small GTPase Rab34 and lysosome-localized RILP^25,26^. Unconventional secretion from autophagosomes during starvation is facilitated by a physical interaction between GRASP55 with LC3-II and LAMP2 ^27^. The generation and maturation of presynaptic vesicle precursors require cross-talk between the small GTPases Rab2 and Arl8B, localized to the Golgi apparatus and lysosomes, respectively ^28^. Post-Golgi transport carriers also shuttle lysosomal components in the axon bidirectionally to promote lysosome maturation ^29^. However, a functional physical coordination between Golgi apparatus and lysosomal membranes has not yet been established.

Skin epidermis constitutes of multiple keratinocyte layers arranged from proliferating to gradually differentiating upper layers, with concomitant changes in intracellular organelles ^30,31^. In the late-stage differentiation, keratinocytes loose conventional intracellular organelles including nuclei, while generating specialized organelles that support epidermis function and homeostasis ^32,33^. Recent reports highlight critical contribution of lysosomes in organelle removal through increased autophagy, which possibly also sustain the nutrient deficient environment in the upper epidermis ^32,34–36^. Previously we demonstrated increased biogenesis and peripheral distribution of lysosomes, and redistribution of the Golgi apparatus during *in vitro* differentiation of human primary keratinocytes. The biogenesis of keratinocyte lysosomes does not follow a conventional mTOR-regulated TFEB/TFE3 pathway. Instead, it relies on an ER stress regulated UPR pathway ^37^. We hypothesized that biochemical alterations in lysosomes are correlated with the changes in keratinocyte physiology during the process of differentiation.

In this study, we show that the lysosomes and dispersed Golgi stacks in differentiated keratinocytes are organized in close physical proximity, and that the lysosomes receive Golgi-derived cargoes that are likely required to maintain functional lysosomes. The close apposition of Golgi lysosomal membranes is specifically mediated by the Golgi tethering protein GRASP65, which associates directly with the lysosome membrane in differentiated keratinocytes. Accordingly, siRNA-mediated depletion of GRASP65 results in the loss of Golgi-lysosome contacts and in the loss of normal lysosomal morphology and function. We further show that the *trans*-Golgi enzyme galactosyl transferase (galT) and secretory cargoes extensively accumulate in the lumen of the lysosomes and are secreted to the cell surface in differentiated keratinocytes, demonstrating a direct contribution of Golgi-derived material to lysosome specialization during keratinocyte differentiation. Finally, we demonstrate that the maintenance of keratinocyte specialized lysosomes critically depends on Golgi function. This study demonstrates that direct contacts with the Golgi apparatus contributes to the biogenesis of specialized lysosomes in differentiated keratinocytes.

## Results

### Lysosomes of differentiated keratinocytes possess conventional lysosomal characteristics

Increased lysosome biogenesis accompanies high calcium-induced differentiation of human primary keratinocytes *in vitro* ^37^. We first compared the lysosomes of proliferative and differentiated keratinocytes (referred to here also as ‘proli.’ and ‘diff.’ respectively) by immunofluorescence microscopy (IFM). IFM analysis for the late endosomal/ lysosomal integral membrane protein LAMP1 revealed that whereas most LAMP1-containing compartments were perinuclear in proliferative cells, they were largely distributed throughout the cytoplasm of differentiated cells (Fig. 1A). Moreover, the average number (lysosomes/µm^2^ area) and size (in μm) of LAMP1-containing lysosomes were significantly higher in differentiated keratinocytes than in proliferative cells (fold increase in lysosome number = 3.63, Fig.1B; fold increase in lysosome size =1.33, Fig. 1C). By Airyscan super-resolution microscopy, the LAMP1-containing lysosomes of differentiated keratinocytes distributed among two different morphologies: bright filled structures and less intense doughnut-shaped structures (represented here as ‘a’ and ‘b’ in the zoomed inset of differentiated cells; Fig. 1A). The LAMP1 intensity of ‘a’ structures was significantly higher than the ‘b’ population (Fig. 1D), but nevertheless we did not observe any notable functional differences between these structures as described further below. Besides ‘a’ and ‘b’, a population of tiny LAMP1 structures was also present, which likely represents post-Golgi LAMP1 carriers (∼100-300nm in diameter) ^38^ (white arrowheads in Fig. 1A). We frequently observed the assembly/fusion of these tiny structures with the larger LAMP1-containing lysosomes in live cell imaging experiments (arrowheads in Fig. S1A and Video S1). By contrast, the LAMP1-containing lysosome population of proliferative keratinocytes mainly appeared as filled and bright structures, similar to the ‘a’ population in differentiated cells. In both proliferative and differentiated keratinocytes, the entire population of LAMP1-positive lysosomes was also positive for the lysosomal membrane proteins CD63 and LAMP2, regardless of their morphology (Figs. 1E and S1B).

**Figure 1.**
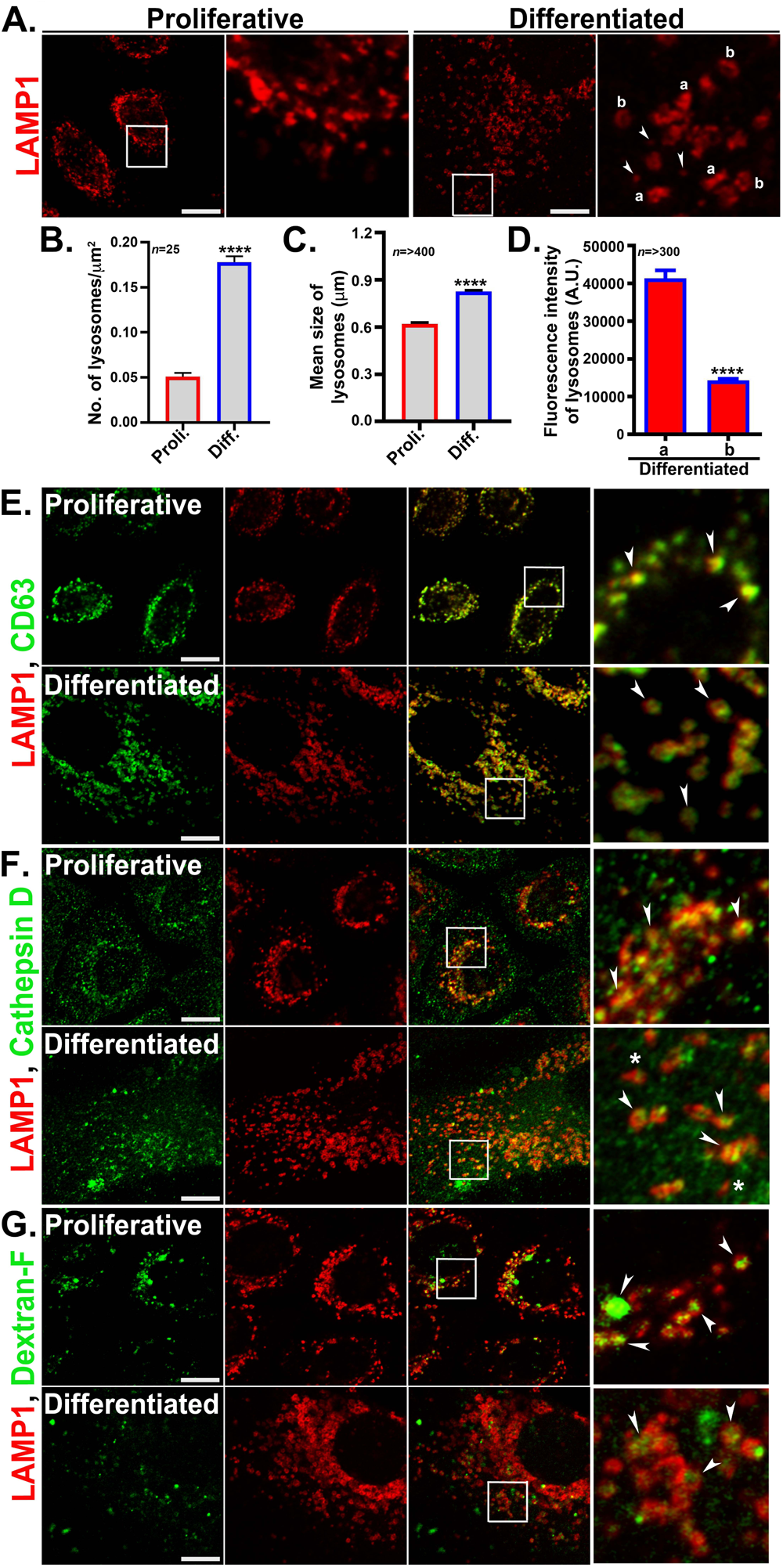
Characterization of the lysosomes in differentiated keratinocytes. (**A**) IFM analysis of the lysosomes of LAMP1 immuno-stained proliferative and differentiated keratinocytes. Two different morphologies of lysosomes ‘a’ and ‘b’ were observed in differentiated keratinocytes. Arrowheads point to the possible LAMP1 carriers. (**B**) Significant increase in lysosome number/μm^2^ cell area in differentiated keratinocytes; n=number of cells. (**C**) Average lysosome size (in μm) is significantly increased in differentiated cells compared to proliferative cells; n=number of lysosomes. (**D**) The graph represents LAMP1 fluorescence intensity of ‘a’ and ‘b’ type lysosomes in differentiated keratinocytes; n=number of lysosomes. (**E**-**G**) IFM analysis of proliferative and differentiated keratinocytes that are co-immunostained for LAMP1 and CD63 (E), cathepsin D (F), or internalized dextran-fluorescein (G) and imaged by Airyscan super-resolution microscope. White arrows in the zoomed boxes point to the colocalization between the markers. White stars in panel F point cathepsin D negative LAMP1 compartments. Scale bar=10 μm. In all images the insets represent 2.5 times magnified white boxed regions.

Next, we tested the functional characteristics of the lysosomes based on the presence of hydrolase cathepsin D which marks the active lysosomes ^39^, and the uptake of the soluble substrates LysoTracker Red DND99 (LR) and dextran-fluorescein. Coimmunostaining experiments revealed that most of the LAMP1-positive lysosomes were positive for cathepsin D in both proliferative and differentiated cells (Fig. 1F, arrowheads). This result was further corroborated by labeling with the cathepsin B substrate magic red (data not shown). LR is a membrane permeable weak base and used as an acidotropic probe to mark acidic compartments. Uptake of LR followed by LAMP1 immunostaining revealed that the lysosomes in proliferative and differentiated keratinocytes were predominantly positive for LR (Fig. S1C). However, a small population of cathepsin D-negative, LR-negative small LAMP1 structures in the peripheral region was also present in differentiated keratinocytes, possibly representing inactive lysosomes (white stars in Fig.1F and Fig. S1C) ^39–41^. Dextran-fluorescein (70 KDa) is a soluble cargo that is internalized by cells and accumulates in the terminal endocytic compartments or lysosomes. Uptake of dextran followed by LAMP1 co-staining revealed a significant dextran accumulation in proliferative keratinocytes, indicating that most LAMP1-containing compartments were accessible to the endocytic pathway. However, dextran accumulation in differentiated keratinocytes was not homogenous (Fig. 1G). This might reflect either a lower rate of internalization in differentiated cells relative to proliferative keratinocytes, a slower rate of endocytic maturation, or a population of lysosomes that do not fuse with late endosomes. Taken together, the above results indicate that the lysosomes of differentiated keratinocytes possess molecular properties of acidic and active conventional lysosomes.

### The Golgi apparatus disassembles and redistributes in close apposition with the lysosomes in differentiated keratinocytes

We next assessed the morphology of the Golgi apparatus in proliferative and differentiated keratinocytes by IFM for GM130 and p230, peripheral membrane proteins of the *cis* and *trans* Golgi, respectively. In proliferative cells, the Golgi had a typical perinuclear arrangement. By contrast, the Golgi in differentiated keratinocytes was highly dispersed (Fig. 2A). Consistently, conventional EM (not shown) and EM analysis of ultrathin cryosections revealed that whereas proliferative keratinocytes harbored a compact Golgi structure with a few (4-10 characteristic buds and vesicles, the Golgi apparatus in differentiated keratinocytes consisted of many short, dispersed ministacks associated with many more 10-100 associated vesicles (Fig. S2G). Despite the dispersion, the association between the *cis*-Golgi marker GM130 and *trans*-Golgi marker p230 (Figs. 2A & 2C first panel) was similar in proliferative and differentiated keratinocytes, consistent with the maintenance of a stacked, polarized Golgi apparatus in the differentiated cells. Golgi dispersion in mammalian cells has been previously associated with a higher Golgi function ^42^. We assessed the binding of Alexafluor-594-labeled wheat germ agglutinin (WGA), a lectin that binds to sialic acid (SA) and N-acetylglucosamine (GlcNAc) on Golgi-modified glycoproteins at the plasma membrane (PM)^42^. WGA binding to the PM was similar in both proliferative and differentiated keratinocytes and was inhibited by treatment with brefeldin A, a small molecule that inhibits Golgi function, indicating that the dispersed Golgi of differentiated keratinocytes was functional (Figs. S2B & S6G). Next, Golgi-mediated secretion was quantitatively measured by the SUnSET assay (SUrface SEnsing of Translation), which measures total protein synthesis and secretion in a given time point ^43,44^. Puromycin is a structural analogue of aminoacyl-transfer RNA that gets incorporated in elongating peptide chains and can be detected by immunoblot using *anti*-puromycin antibody. Consistently, no puromycin signal was detected in cells treated with cycloheximide (CHX) that inhibits new protein synthesis (Fig. 2B). Remarkably, total protein secretion was significantly higher (∼2.5 fold) in differentiated keratinocytes compared to the proliferative cells (Figs. 2B and S2A). On the other hand, protein secretion was strongly inhibited by Golgicide-A, a functional inhibitor of the Golgi apparatus (Fig. 2B) (see also Fig. 6F). Collectively, these data suggest that Golgi dispersion during keratinocyte differentiation correlates with increased secretion.

**Figure 2.**
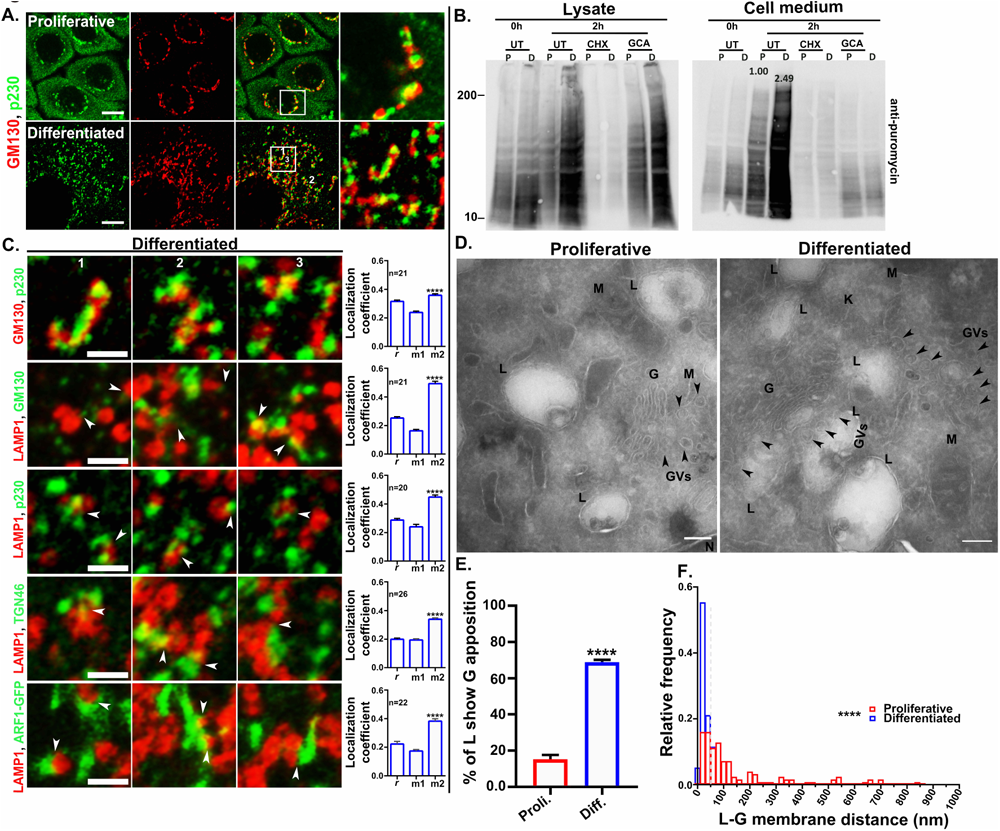
Golgi stacks and lysosomes are closely apposed in differentiated keratinocytes. **(A)** IFM analysis of proliferative and differentiated keratinocytes co-stained for the Golgi markers GM130 and p230. (**B**) Total protein synthesis and secretion from the proliferative and differentiated keratinocytes was measured by the SUnSET assay. Total protein secretion at 0h and 2h were measured with or without the presence of small molecules. P= proliferative and D = differentiated conditions. UT= untreated, CHX= cycloheximide, GCA= Golgicide-A. Total synthesis and secretion is represented as lysate and in cell medium/supernatant respectively. (**C**) co-immunostaining of the representative lysosome marker LAMP1 with the Golgi markers GM130 (representing *cis*/medial Golgi), p230 (*trans* Golgi), TGN46 (*trans* Golgi network) or expressed ARF1-GFP (ubiquitous in Golgi). Cells were visualized by Airyscan super-resolution microscope. Represented insets are the 2.5 times magnified numbered (1-3) regions of the cells shown in Figs. 2A & S2E. Arrowheads represent the close apposition/contact between Golgi and lysosomes. Scale bar=2.5 μm. Pearson’s correlation coefficient (*r*) and Mander’s overlap coefficients (M1 and M2) were calculated in differentiated cells from at least 3 different experiments. n=number of cells. **** on the graphs indicates a significant difference between M1 and M2 values. **(D)** Ultrathin cryosections of proliferative and differentiated keratinocytes were analyzed under TEM. Arrowheads represent the Golgi overlap to the lysosomes. Golgi (G); lysosome (L); mitochondria (M); keratin fibers (K) and, nucleus (N). Scale bar=200 nm. (**E**) The graph is representing % of lysosomes (L) that are closely apposed (within 100 nm) to the Golgi. (**F**) The distance (nm) between lysosomes (L) from the closest Golgi membrane in proliferative and differentiated cells. Dotted grey line on the graph was drawn on 50 nm on the X axis as a reference. The value of significance was calculated by Mann-Whitney non-parametric test. **** = *P* ≤ 0.0001.

The distribution of the dispersed Golgi elements in differentiated keratinocytes appeared roughly similar to that of the lysosomes in these cells. To test if the dispersed Golgi elements were associated with lysosomes, we performed IFM for LAMP1 relative to markers of distinct Golgi regions. Interestingly, labeling for each of the tested Golgi markers was closely apposed to labeling for LAMP1 (Fig. 2C, white arrowheads; Fig. S2E). Organelles such as early endosomes (labeled with EEA1) and mitochondria (labeled with Mito tracker red) did not show a similar apposition to LAMP1 inspite of their discrete distribution in the cytosol of differentiated keratinocytes (Figs. S2C & S2D). Partial overlap between LAMP1 and Golgi markers was quantified by Mander’s overlap coefficient. We considered GM130 and p230 overlap as a suitable reference for this quantification since GM130 and p230 partially overlap inspite of their localization onto the same organelle, the Golgi apparatus (Fig. 2A, 2C first panel). The association between LAMP1-LAMP2 and LAMP1-DAPI was respectively used as positive and negative controls for the quantification method (Fig. S2H; positive control: *r*=0.86±0.01, M1=0.82±0.02 and M2=0.85±0.02); negative control: *r*=0.04±0.01, M1=0.01±0.00, and M2=0.02±0.01). In our quantifications, Mander’s overlap coefficient M1 represents overlap of LAMP1 with the Golgi markers, and M2 values represent Golgi markers overlap with LAMP1. As shown in Fig. 2C, labeling for GM130 overlapped as well with labeling for lysosomes (white arrows; *r*=0.25±0.01, M1=0.16±0.01 and M2=0.49±0.02) as with labeling for p230 (*r*=0.31±0.01, M1=0.24±0.01 and M2=0.36±0.01). Accordingly, p230 labeling overlapped with LAMP1 as well as GM130 (Fig. 2C; *r*=0.29±0.01, M1=0.24±0.02, and M2=0.45±0.02). Although the correlation between labeling for either TGN46 or ARF1-GFP and LAMP1 was lower than that observed with GM130 and p230, their overlap (M2) with the lysosomes was comparable to other Golgi markers (Fig. 2C; TGN46: *r*=0.20±0.01, M1=0.19±0.01, and M2=0.34±0.01; ARF1-GFP: *r*=0.22±0.02, M1=0.17±0.01, and M2=0.38±0.02). Similar quantification in proliferative keratinocytes also showed high overlap between LAMP1 and Golgi markers which was merely due to their perinuclear restricted distribution and was not specific, since all the tested organelles showed similar pattern of association including mitochondria and early endosomes (Figs. S2 C, D & E). In conclusion, Golgi proteins showed a higher overlap tendency with the lysosomes (but not vice versa) in differentiated keratinocytes (refer to M1 and M2 values in respective graphs, Fig. 2C).

To better understand the association between Golgi and lysosomes in differentiated keratinocytes, we performed electron microscopy analyses. Surprisingly, unlike conventional lysosomes in other cell types that display an electron dense and/or multilaminar appearance ^45^, keratinocyte lysosomes predominantly displayed a vacuolar morphology (Fig. 2D). Therefore, we first defined the lysosomes by LAMP1 immunogold labeling of ultrathin cryo-sections (Fig. S2F). We considered a double membraned organelle as a lysosome by the presence of at least two gold particles (Fig. S2F). TEM analysis of the ultrathin cryo-sections of differentiated keratinocytes revealed that a significant population of lysosomes were closely apposed to Golgi ministacks (Fig. 2D, right panel, black arrowheads). Quantification revealed that 69 ± 2.67% of the lysosomes in differentiated keratinocytes were within 100 nm of a Golgi stack, compared to only 15%± 2.67 in proliferative keratinocytes (Fig. 2E). Further, quantification of the distance (nm) between the lysosome membrane and the nearest Golgi membranes revealed a regular distribution between these organelles (within ≤ 50 nm) in differentiated keratinocytes, compared to a random and wide distribution in proliferative keratinocytes (Fig. 2F). Collectively, these results confirm that lysosomes in differentiated keratinocytes are in close physical proximity to Golgi membranes in differentiated keratinocytes.

### The peripheral Golgi protein GRASP65 surrounds lysosomes in differentiated keratinocytes

GRASP65 and GRASP55 are peripheral Golgi membrane proteins that regulate Golgi ribbon morphology through oligomerization ^9,17^. We therefore hypothesized that the Golgi dispersion during keratinocyte differentiation was due to impairment of either GRASP55 or GRASP65 function. Initial studies showed that GRASP55 or GRASP65 mRNA expression levels were not significantly down-regulated during keratinocyte differentiation (Fig. S3B). Moreover, overexpression of GRASP65-GFP, GRASP55-mCherry, or both in differentiated keratinocytes could not reverse Golgi dispersal (Fig. S3A); co-expressed GRASP55 and GRASP65 fusion proteins both localized to the dispersed Golgi stacks (Fig. S3A), and the pattern of Golgi dispersion was similar to the endogenous pattern of Golgi in differentiated keratinocytes (Fig. S3H), suggesting that the GRASPs associate passively with the dispersed Golgi elements.

In addition to the prominent localization to dispersed Golgi stacks and a diffuse cytoplasmic cohort, GRASP65-GFP also appeared in unique ring like structures (Fig. 3A, arrows, and arrowheads). These GRASP65-GFP-containing ring-like structures did not label for GRASP55-mCherry (Fig. S3A) or for other Golgi proteins such as ARF1 (data not shown), and were detected by live cell imaging in ∼95% of the differentiated keratinocytes that expressed GRASP65-GFP but only ∼9% of corresponding proliferative keratinocytes (with 1-2 rings/cell) (Figs. 3B and S3C& D; Video S3). The GRASP65-GFP ring like structures were best visible by live cell imaging; they were less detectable in fixed cells (observed in ∼60% of the cells) and not detectable by IFM for endogenous GRASP65 (Figs. 3B and S3H), suggesting that they are likely sensitive to fixation.

**Figure 3.**
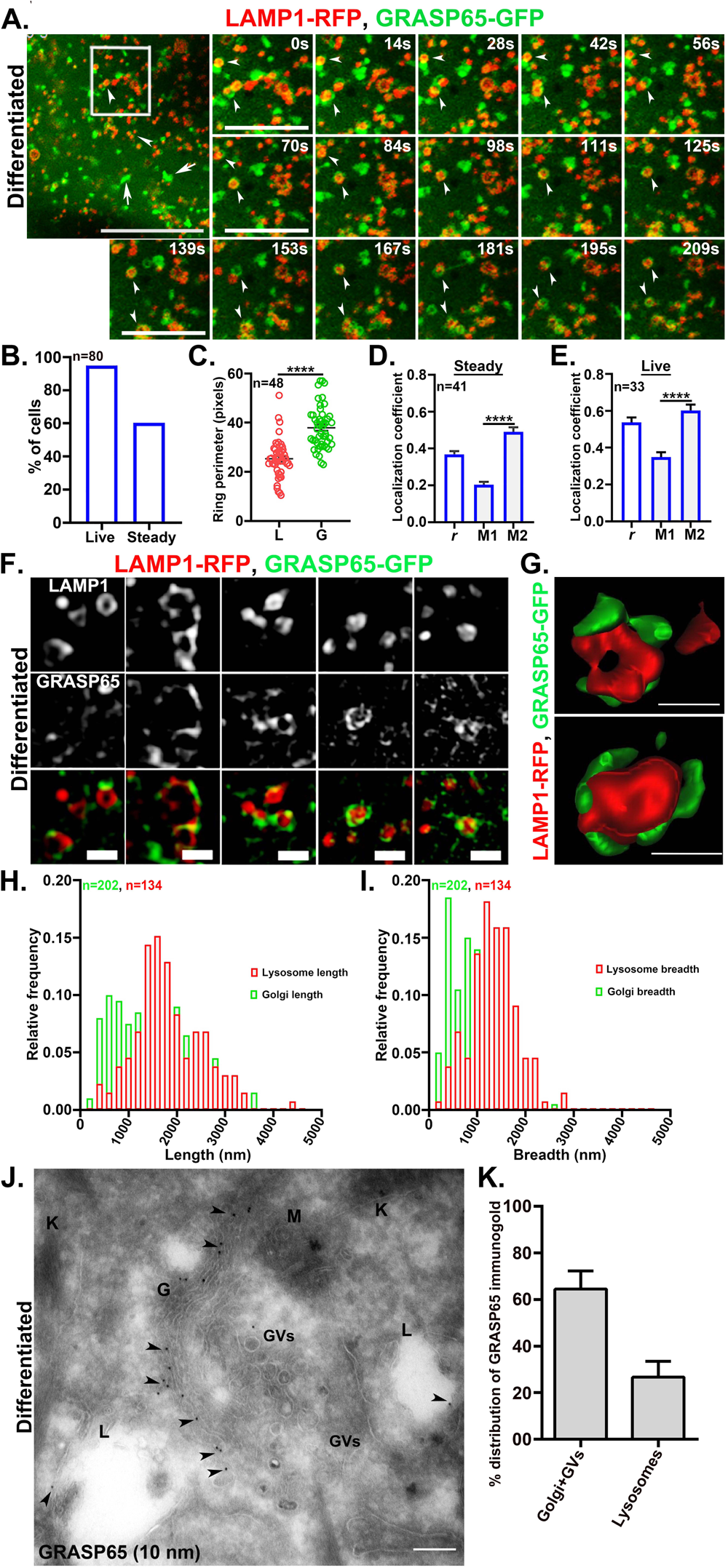
The peripheral Golgi protein GRASP65 surrounds the lysosomes of differentiated keratinocytes. (**A**) Live cell imaging of differentiated keratinocytes expressing GRASP65-GFP and LAMP1-RFP by Airyscan super-resolution microscope. The arrows and arrowheads represent respectively to the Golgi stacks and GRASP65-GFP ring-like structures. Scale bar=5 μm. Time series of the white-boxed region are presented. Arrowheads represent the dynamics of GRASP65-GFP ring-like structures with respect to the lysosomes. Scale bar=2.5 μm. (**B**) The graph represents % of cells positive for GRASP65-GFP ring-like structures; n = number of cells. (**C**) The perimeter of the lysosomes (L) and, perimeter of the respective GRASP65 ring-like structures (G) was measured; n=number of lysosomes surrounded by prominent GRASP65 ring structures. (**D**-**E**) Pearson’s correlation coefficient (*r)* and Mander’s overlap coefficients (M1 and M2) in steady and live conditions were calculated and plotted as mean ±s.e.m.; n=number of cells. (**F**) SIM analysis of the GRASP65-GFP and LAMP1-RFP association in differentiated keratinocytes. Scale bar=1 μm. (**G**) analysis of GRASP association on the lysosomes by 3D rendering. Scale bar=500 nm. (**H-I**) The length and breadth analysis (values in nm) of lysosomes (LAMP1-RFP) and Golgi (GRASP65-GFP) were plotted as a function of fractions. ‘n’ in green=number of GRASP65-positive Golgi structures; ‘n’ in red=number of LAMP1-positive lysosome structures. (**J**) Ultrathin cryosections of differentiated keratinocytes were immunolabelled for GRASP65 (10 nm gold particles) and imaged under TEM. Arrowheads represent the localization of GRASP65 to the Golgi and lysosomes. Abbreviations are Golgi (G); lysosome (L); mitochondria (M); keratin fibers (K) and Golgi vesicles (GVs). Scale bar=200 nm. (**K**) Quantification of the % distribution of GRASP65 immunogold beads in Golgi and lysosomes. Quantification was performed with the images of two independent experiments.

Interestingly, the GRASP65-GFP ring like structures surrounded the lysosomes of differentiated keratinocytes, as observed upon co-expression with LAMP1-RFP (Fig. 3A, arrowheads; Video S2). Accordingly, the perimeter of the GRASP65-GFP rings (denoted as G; 37.9±1.3 pixels) was significantly larger than that of the LAMP1-RFP rings (denoted as L; 25.3±1.1 pixels) (Fig. 3C). This special association of GRASP65 to the lysosomes in differentiated keratinocytes also reflected by highest value of colocalization (‘r’) among the Golgi proteins tested, and by the fact that it was sensitive to fixation (*r*=0.54±0.03 in live; *r*=0.39±0.02 in steady Figs. 3E and 3D). Accordingly, in proliferative keratinocytes, Pearson’s correlation coefficient remained unaltered between live and steady states (*r*=0.47±0.03 in live and 0.44±0.02 in steady; Figs. S3F and S3E). Because GRASP55 has been implicated in autophagosome fusion and secretion ^27,46,47^, we tested whether the LAMP1-positive structures are autolysosomes by co-immunostaining for LAMP1 and LC3. LC3 did not significantly overlap with LAMP1-positive structures in either proliferative or differentiated keratinocytes (Fig. S3G, arrowheads), indicating that these structures are not autolysosomes. Overall, these studies revealed that Golgi peripheral protein GRASP65 surrounds the lysosomes of differentiated keratinocytes.

To better appreciate the spatial relationship between GRASP65-GFP and the lysosomes in differentiated keratinocytes, we analyzed GRASP65-GFP-and LAMP1-RFP-expressing cells by structured illumination microscopy (SIM) that gives a better resolution in the axial plane than Airyscan super resolution (in Airyscan: XY=∼120nm, Z=∼350nm; in SIM: XY=∼100nm, Z= ∼250nm). Whereas GRASP65-GFP appears as complete rings around lysosomes by Airyscan super resolution microscopy (Fig. 3A), GRASP65-GFP appeared as discontinuous patches arranged around the periphery of the lysosomes in SIM imaging, particularly upon the 3D rendering of the images (Figs. 3F and 3G). The average number of GRASP65 patches per lysosome ranged between 1 to 3. Further, the length and breadth values of the GRASP65 positive Golgi structures and LAMP1-positive lysosome structures were distributed independently (Figs. 3H and 3I), which suggests that GRASP65 association with the lysosomes is an active association and merely not due to their structural similarity (Fig. 3F).

Immunogold labeling of ultrathin cryosections of differentiated keratinocytes also revealed the presence of GRASP65 on the lysosome membrane (Fig. 3J). As expected, the majority of the GRASP65 immunogold was associated with the Golgi stacks and Golgi vesicles (GVs) (64±8%) (Figs. 3J and 3K). However, a significant fraction was also present on the lysosomes (27±7%) (Figs. 3J and 3K); by contrast, GRASP65 labeling in the nucleus and mitochondria were negligible (not shown). Of note, approximately 46% of the total lysosome population was positive for GRASP65 with an average frequency of one GRASP65 immunogold particle per lysosome (Fig. 3J). Nonetheless, endogenous detection of GRASP65 by immunogold labelling can be described by the fact that even single immunogold is detectable by TEM, but a minimum fluorescence intensity over background is required to be detected by IFM. Altogether, these results confirm the presence of GRASP65 on the periphery of lysosomes in differentiated keratinocytes.

### GRASP65 mediates Golgi lysosome apposition in differentiated keratinocytes

Given that GRASP65 uniquely surrounded lysosomes in differentiated keratinocytes, we tested whether GRASP65 was necessary for lysosome-Golgi apposition in these cells by assessing the impact of depleting GRASP65 with siRNA. Cells were treated with siRNA to GRASP65, GRASP55 or a non-targeting siRNA as negative controls, or to GM130 which mediates interaction of GRASP65 with Golgi membranes ^19^. Approximately 70% depletion of GRASP65 and GRASP55 mRNA was achieved with the respective siRNAs (Fig. S4E), and depletion of GRASP65, GRASP55, and GM130 were additionally confirmed by IFM (Figs. S4B, S4C, and S4D) and by immunoblotting for GRASP65 (Fig. S4F). Golgi dispersion in differentiated keratinocytes was not impacted by treatment with any of the siRNAs, indicating that neither GRASP65, GRASP55, nor GM130 are required for Golgi dispersion during differentiation (Fig. 4A, data not shown). However, individual Golgi elements in GRASP65 siRNA-treated cells was significantly bigger in volume compared to control siRNA-treated cells (Fig. S4H).

**Figure 4.**
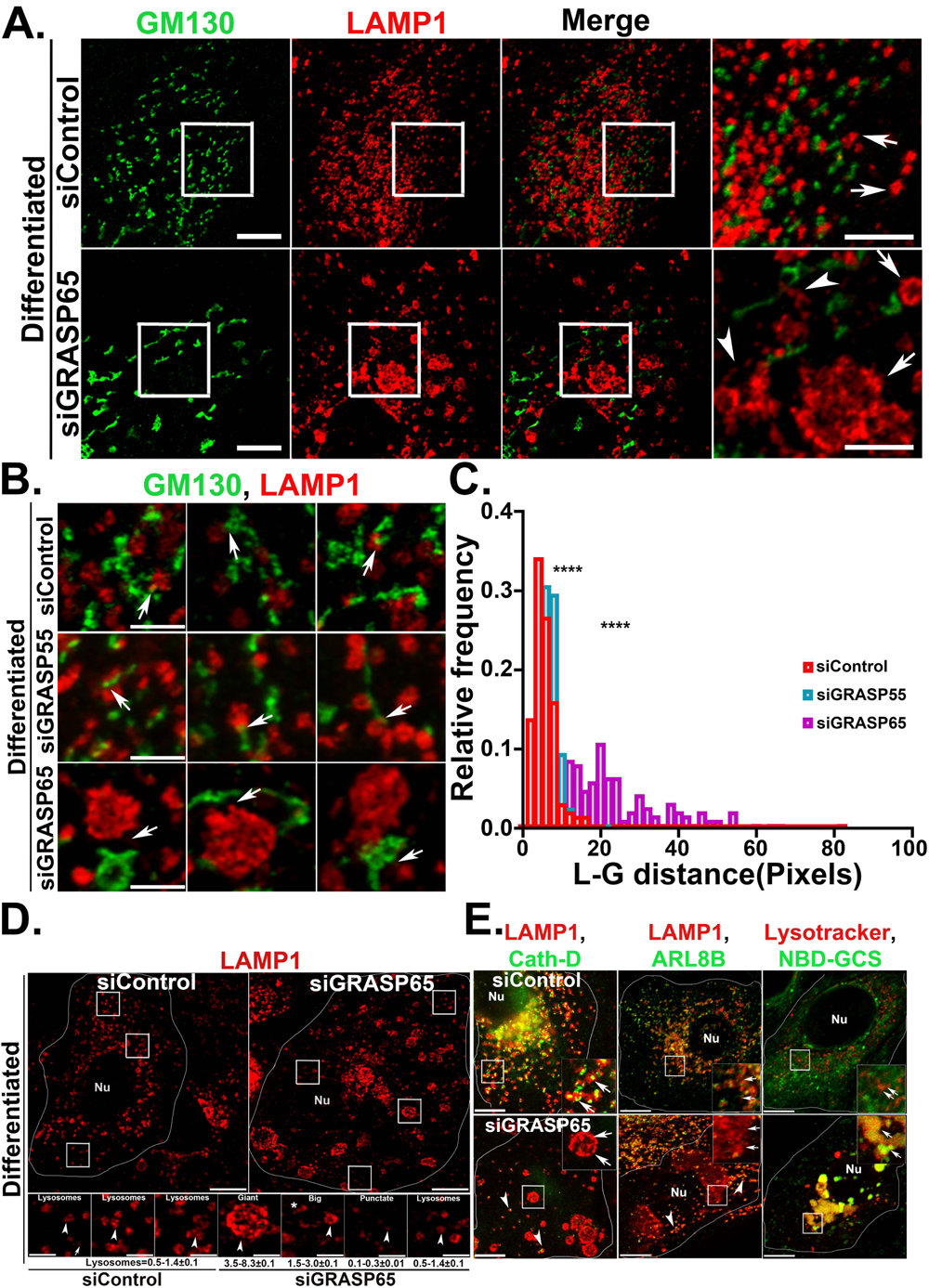
GRASP65 mediates Golgi-lysosome contacts in differentiated keratinocytes. **(A)** IFM analysis of siControl and siGRASP65 treated differentiated keratinocytes co-immunostained for GM130 and LAMP1. Cells were imaged by Airyscan super resolution microscope. Insets represent 2.5 times magnified white boxed regions. Arrows indicating to the lysosome morphology in siControl and siGRASP65 cells. Arrowheads represent the tubular LAMP1 positive structures in siGRASP65 cells. Scale bar=10 μm (main inset) and 5 μm (magnified inset) respectively. (**B)** IFM analysis of siControl, siGRASP55, and siGRASP65 treated cells stained for GM130 and LAMP1. White arrows indicate the loss of Golgi-lysosome apposition in siGRASP65, but not in siControl and siGRASP55 treated cells. (**C**) The distance between lysosome (L) and the closest Golgi stack (G) in respective siRNA treated cells was measured by using Fiji software. The distribution of the values representing G-L distance was plotted as a function of fractions. The value of significance was calculated by Kolmogorov-Smirnov non-parametric analysis. (**D**) Representation of the overall lysosome population in GRASP65 siRNA treated cells compared to control siRNA. The lysosome population according to their morphology and measured size range (in diameter) is presented. Scale bar= 10μm and 2.5 μm respectively. Nu= nucleus. (**E**) A brief functional characterization of the giant LAMP1-positive structures in GRASP65 siRNA treated differentiated keratinocytes. Cells were either co-immunostained for LAMP1with Cathepsin D, Arl8B or, uptake of NBD glucosylceramide (GCS) along with lysotracker red (1^st^, 2^nd^ and 3^rd^ panel). Represented insets are 2.5 times magnified white boxed areas of the respective cells. Arrows represent to the characteristics of the giant LAMP1 positive structures with respect to specific markers. Arrowheads represent regular population of lysosomes in GRASP65 siRNA treated cells. Scale bar= 10μm. Nu= nucleus.

Interestingly, analysis of the distribution of lysosomes relative to the dispersed Golgi elements revealed a prominent loss of Golgi-lysosome apposition in GRASP65 knockdown cells compared to GRASP55 or control siRNA treated cells (Fig. 4B). This was quantified as a significantly increased average distance and a wider and more random distribution of distances between the lysosomes and Golgi stacks in cells depleted of GRASP65 but not GRASP55 (Figs. 4B arrows; Fig. 4C). These data indicate that GRASP65 expression is necessary for the close apposition of Golgi elements and lysosomes in differentiated keratinocytes.

In addition to the reduced apposition to Golgi elements, the lysosomes in GRASP65 siRNA-treated cells were more varied in morphology and size (diameter) than in control siRNA-treated cells. In control siRNA, the LAMP1 positive lysosomes mostly appeared as doughnut shaped structures with a mean size range between 0.5-1.4 μm (Fig. 4D). In contrast, the lysosome population in GRASP65 siRNA cells consisted of prominent “giant” structures (diameter 3.5 - 8.3 μm), comparatively “big” structures relative to lysosomes in control siRNA-treated cells (1.5 -3.0 μm), “regular” structures similar in size to those in control siRNA-treated cells, tiny “punctate” structures (0.1 -0.3 μm), and some tubulated structures (Fig. 4D; arrows in siGRASP65 panel Fig.4A). Among these, LAMP1 positive giant structures were the most prominent characteristic of GRASP65 depletion. The giant LAMP1 structures varied in size, morphology, number (∼4-10 per cell), and distribution to the periphery or perinuclear regions. By contrast, treatment with siRNAs to GRASP55 or GM130 did not show significant alteration in lysosome morphology (Figs. S4C and S4D). Moreover, depletion of GRASP65 in proliferative keratinocytes did not significantly affect lysosome morphology, reinforcing the functional specificity of GRASP65 for lysosomes in differentiated keratinocytes (Fig. S4A). Altogether, these data suggest essential roles of GRASP65 in facilitating Golgi-lysosome apposition in differentiated keratinocytes, the loss of which possibly resulted in the aberration of lysosome morphology.

To determine whether the characteristic giant LAMP1 positive structures of GRASP65 siRNA-treated cells were functional, we performed additional fluorescence microscopy analyses. The giant LAMP1-positive structures accumulate lysotracker red and internalized dextran-fluorescein (Figs. 4E panel3; S4G; not shown), indicating that they are acidic and accessible to the endocytic pathway. However, while regular lysosomes in these cells labeled for cathepsin D and ARL8 (Fig. 4E, first two panels, arrowheads), the giant lysosomes did not (Figs. 4E first two panels, arrows; S4G). Consistent with a defect in lysosomal degradation, exposure of control siRNA-treated cells to the fluorescent lysosomal substrate NBD-glucosylceramide resulted in substantial clearance, whereas GRASP65 siRNA-treated cells massively accumulated the substrate in giant acidic lysosomes that labeled for lysotracker (Figs. 4E panel3; S4G). These results suggest that the giant lysosomes in GRASP65-depleted cells are functionally aberrant. Altogether, these data suggest an active role of GRASP65 in mediating the physical apposition between Golgi and lysosome, and that this apposition is required for the functional maturation/maintenance of lysosomes in differentiated keratinocytes.

### Lysosomes in differentiated keratinocytes accumulate secretory cargoes and a *trans*-Golgi enzyme, and are secretory in nature

To understand the functional significance of Golgi lysosome cross-talk, we examined the fates of Golgi resident enzymes and secretory cargo in differentiated keratinocytes. β-1,4 galactosyltransferase1 (galT) is a *trans*-Golgi resident enzyme which, in some cells, cycles through the TGN and the plasma membrane ^48^. Mannosidase II (ManII) is a medial Golgi enzyme. We assessed the dynamic localization of galT-RFP and ManII-GFP with respect to the LAMP1 positive lysosomes of differentiated keratinocytes. ManII-GFP localized exclusively to the dispersed Golgi stacks in close apposition to the lysosomes (Fig. S5A), consistent with the distribution of GRASP55, GM130, p230, and ARF1-GFP (Fig. 2). By contrast, whereas a cohort of galT-RFP localized to the dispersed Golgi stacks, a major fraction of galT-RFP appeared as bright round structures that localized to the lumen of LAMP1-GFP labeled lysosomes in differentiated keratinocytes (Fig. 5D last panel; Fig. 5A, white arrows and arrowheads). Accordingly, EM analysis of ultrathin cryosections that were immunogold labeled for endogenous galT revealed the presence of galT in ∼47% of the lysosomes in differentiated keratinocytes, with a frequency of 1 to 3 gold particles per lysosome (Fig. 5B). Unlike GRASP65 labelled immunogold, for which only a small cohort localized to lysosomes, immunogold labeling for galT was higher in lysosomes than in the Golgi apparatus and Golgi vesicles (48.3±7.3% in lysosomes and 41.3±2.9% in Golgi/vesicles; negligible labeling was detected in other organelles) (Fig. 5C and Fig. 3K). These results document that a significant fraction of a *trans*-Golgi enzyme is delivered to the lumen of lysosomes in differentiated keratinocytes.

**Figure 5.**
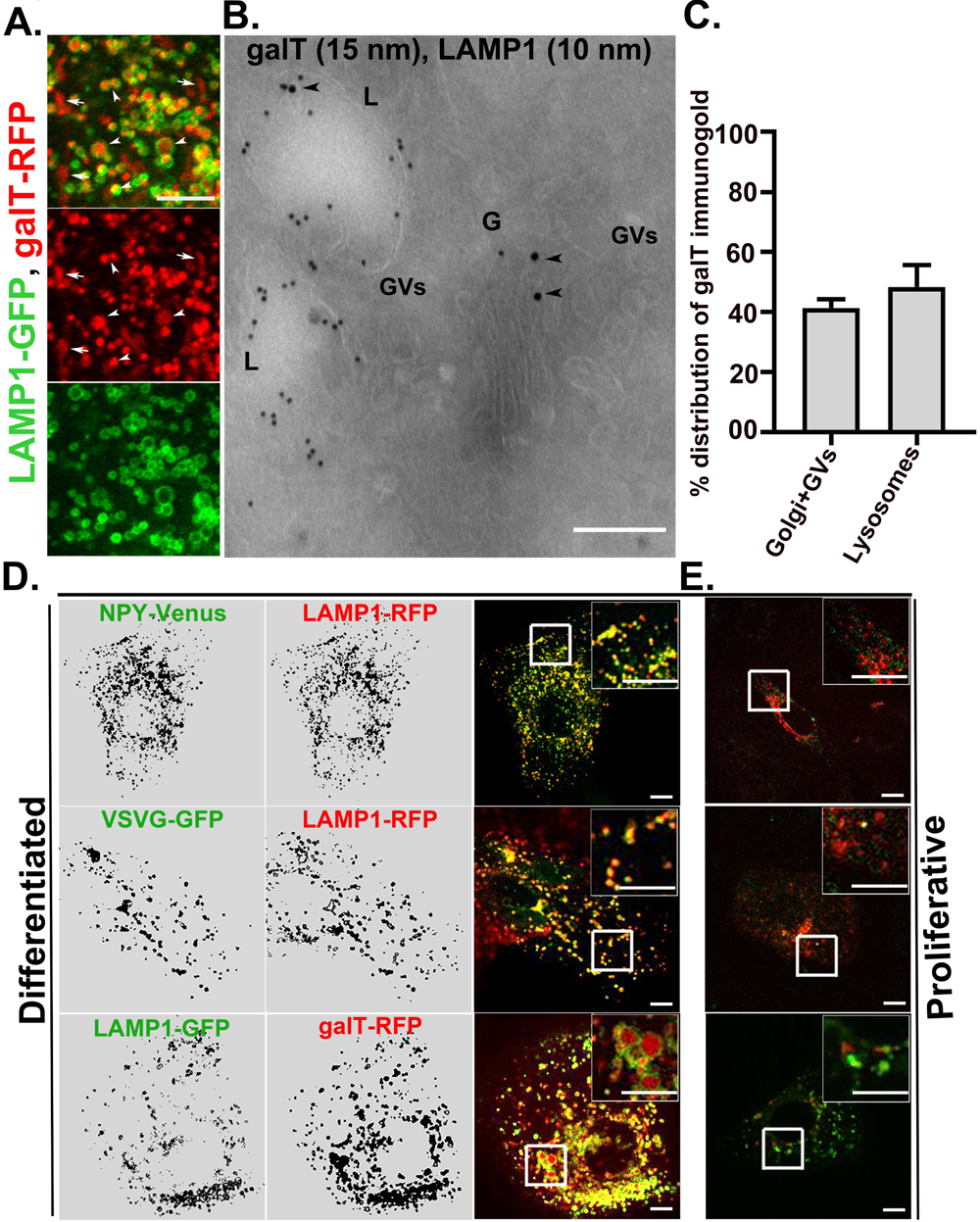
Keratinocyte lysosomes receive secretory cargo. (**A**) Live cell analysis of differentiated keratinocytes expressing galT-RFP and LAMP1-GFP. Imaging was performed by Airyscan super-resolution microscope. Arrows indicate to the Golgi localized galT and arrowheads indicate the presence of galT in the lumen of lysosomes. Scale bar=5 μm. (**B**) Ultrathin cryosections of differentiated keratinocytes were co-immunolabelled for galT (15 nm) and LAMP1 (10 nm). Black arrowheads represent the presence of galT in the Golgi and lysosome. G=Golgi; GVs=Golgi vesicles and L=lysosome. Scale bar=200 nm. (**C**) Quantification of the % of galT immunogold on the lysosomes and Golgi. The average percentage of the distribution of galT immunogold was calculated from three independent imaging experiments. (**D-E**) localization of the *trans* Golgi cargo and secretory cargo NPY-venus and VSVG-GFP in differentiated (D) and in proliferative keratinocytes (E). Cells were co-expressed with indicated constructs and live cell imaging was performed by a spinning disk microscope. White boxed region in the merged panel was magnified 2.5 times. Scale bar=10 μm.

To assess the fate of secretory cargoes in differentiated keratinocytes, we analyzed the distribution of ectopically expressed fluorescent-tagged neuropeptide Y (NPY) and vesicular stomatitis virus glycoprotein (VSVG). Golgi to PM transport of NPY and VSVG is mediated by Rab6-positive *trans-*Golgi-derived secretory carriers ^49,50^. Strikingly, we observed selective and extensive accumulation of both NPY and VSVG within the lysosomes of differentiated keratinocytes (Fig. 5D, panels 1 and 2). The extent of localization of NPY or VSVG with the lysosomes was comparable to that of secretory/exocytic vesicles as described in other cell types ^49^. By contrast, in proliferative keratinocytes, a very small fraction of lysosomes appeared positive for NPY, VSVG or galT (∼ 14% of cells) (Fig. 5E). Further, we used the temperature sensitive mutant of VSVG (ts045VSVG-GFP) co-expressed with lysosome marker LAMP1-RFP to assess secretory properties of keratinocyte lysosomes. The ts045VSVG misfolds and is retained in the ER at 40^0^C and moves out of the ER to Golgi and gradually to the cell surface at a permissive temperature of 32^0^C (Yoshino et al., 2005). As shown in Fig. S5B, at 40^0^C, ts045VSVG-GFP appeared in the ER with a reticular pattern which does not co-localize with LAMP1 positive lysosomes. After 30 minutes incubation at 32^0^C, the ts045VSVG signal was extensively localized with the LAMP1 labeled lysosomes, with a similar extent to the full length VSVG-EGFP (Fig. 5D). With increased incubation at 32^0^C for 60 minutes and 90 minutes, increasing signal of ts045VSVG and LAMP1 were appeared in the cell surface (Fig. S5B), suggesting that secretory cargo containing lysosomes in differentiated keratinocytes secrete/exocytose to the cell surface. It was previously shown that the lysosomal acidifying inhibitor bafilomycin A1 inhibits lysosome exocytosis ^51^. Accordingly, in the presence of bafilomycin A1, the total protein secretion was inhibited by ∼70% in differentiated keratinocytes as measured by the SUnSET assay (Fig. S5C). The inhibition in total protein secretion which is presumably due to the inhibition in lysosome exocytosis, correspond well with the perinuclear accumulation of lysosomes in differentiated keratinocytes (Fig. S5D). Taken together, these results suggest that lysosomes in differentiated keratinocytes receive direct input from the Golgi and contribute to the total protein secretion from differentiated keratinocytes. We refer to these lysosomes as ‘specialized’ because they have both degradative and secretory functions.

### A functional Golgi is essential for lysosome maintenance in differentiated keratinocytes

To directly visualize Golgi transport intermediates contacting lysosomes in differentiated keratinocytes, we used live cell imaging to analyze cells co-expressing GFP-tagged ARF1 with LAMP1-RFP. Interestingly, ARF1-GFP localized to the dispersed Golgi stacks and to unique tubulo-vesicular structures (Fig. 6A). The ARF1-containing tubulo-vesicular dynamic structures were visible both in live imaging and in fixed cells, and were frequently seen contacting/fusing with lysosomes in time lapse experiments (Fig. 6A, arrowheads; Video S4, Figs. 6B & 6C). A few small tubulo-vesicular structures were also observed in proliferative keratinocytes (arrowheads Figs. S6A, Video S5 and, S6B) but were not detected in other cell types such as HeLa (Fig. S6F). GFP alone or fused to other Golgi-associated proteins did not generate similar structures, confirming that ARF1 specifically associated with these dynamic tubulo-vesicular compartments in keratinocytes. To determine whether ARF1 regulates lysosome dynamics, the GFP-tagged dominant negative mutant of ARF1 (ARF1^T31N^-GFP) was co-expressed with LAMP1-RFP. As expected, ARF1^T31N^-GFP localized exclusively to the cytosol (Fig. 6D), and resulted in disruption of the Golgi to small punctate structure in the perinuclear region (Fig. S6C). Interestingly, ARF1^T31N^-GFP expression in differentiated keratinocytes resulted in the complete loss of LAMP1 positive lysosomes, leaving only very small LAMP1-positive punctate structures accumulated in the perinuclear area (Fig. 6D, arrows, and arrowheads; size range 0.1 -0.3 μm in diameter). ARF1^T31N^-GFP expression did not significantly affect lysosome distribution in proliferative keratinocytes (Fig. S6D) or HeLa cells (Fig. S6F). These data indicate that ARF1 function is specifically required for lysosomal maintenance in differentiated keratinocytes.

**Figure 6.**
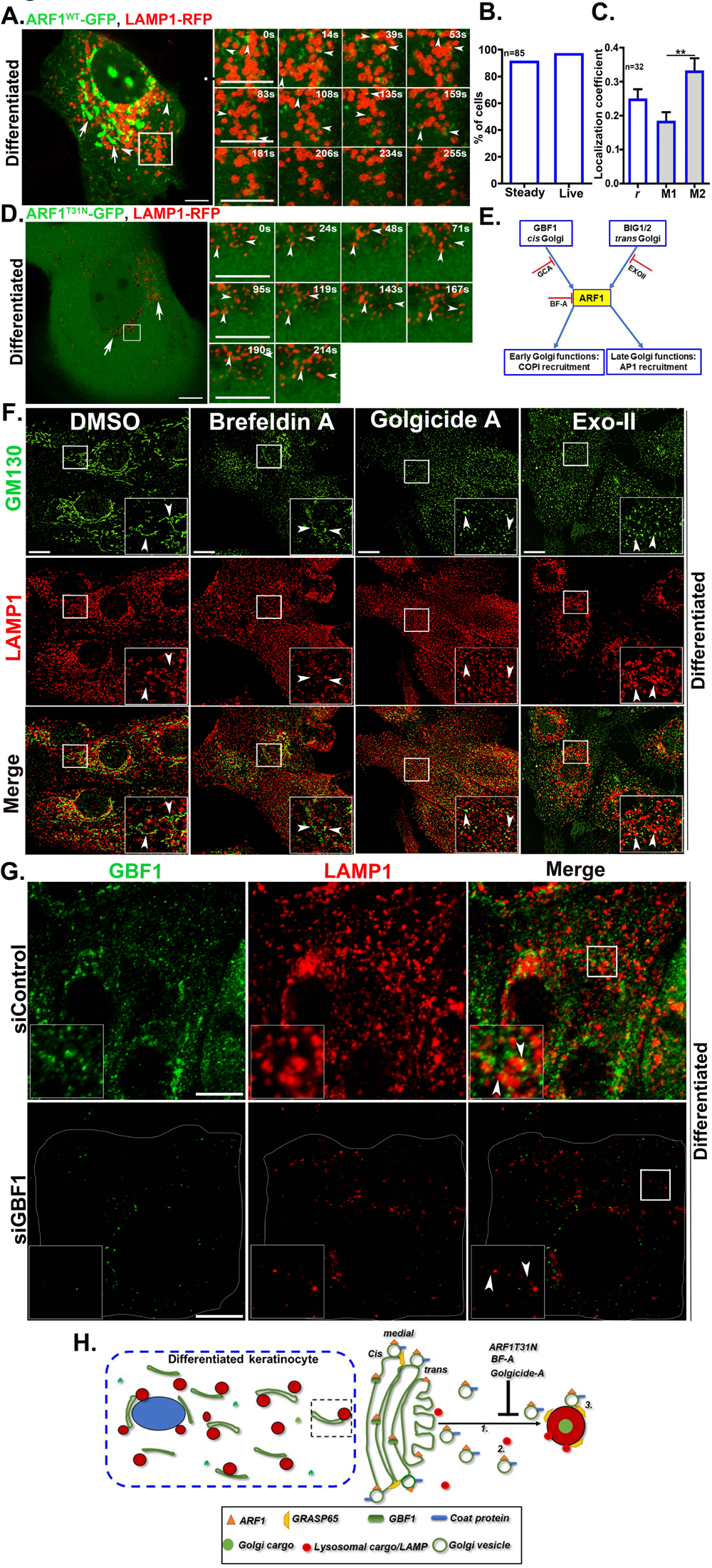
Maintenance of keratinocyte lysosomes is dependent on the Golgi function. (**A**) Time lapse imaging of differentiated keratinocytes expressing ARF1^WT^-GFP and LAMP1-RFP. Arrows and arrowheads respectively represent the localization of ARF1-GFP to the Golgi stacks and to small tubulo-vesicular structures. Time series of the white boxed region are presented. Scale bars=10μm (in the main inset) and 2.5μm (time series). (**B**) The graph represents % of cells positive for tubulo-vesicular structures upon ARF1^WT^-GFP expression in live and steady states. (**C**) Localization coefficients (Pearson’s correlation coefficient *r* and, Mander’s overlap coefficients M1 and M2) were calculated and plotted as mean±s.e.m. ns=non-significant. (**D**) Time series imaging of differentiated keratinocytes expressing ARF1^T31N^-GFP and LAMP1-RFP. Arrows indicate to the cytosolic localization of ARF1^T31N^-GFP and arrowheads (in the insets) point to the punctate LAMP1 structures in the perinuclear region. Scale bar=10μm (in the main inset) and 2.5μm (time series). (**E**) Schematic representation of ARF1 and its GEFs at *cis* and *trans* Golgi, and their respective pharmacological inhibitors. (**F**) Keratinocytes treated with respective pharmacological inhibitors were co-immunostained for GM130 and LAMP1. Insets represent 2.5 times magnification of the white boxed region. Arrowheads indicate to the Golgi with respect to lysosomes in all treatment conditions. n=number of cells. Scale bar=10 μm. (**G**) Disappearance of lysosomes upon siRNA mediated inhibition of GBF1 (**H**) Proposed model describes the generation/maintenance of specialized secretory lysosomes in differentiated keratinocytes through constant Golgi input (lysosomal/Golgi cargoes) likely to be mediated by Golgi derived vesicles. GBF1 mediated ARF1 activation is required for the generation of Golgi vesicles. GRASP65 facilitates receiving of Golgi input/secretory vesicles through Golgi-lysosome apposition in differentiated keratinocytes.

ARF1 is activated by several guanine nucleotide exchange factors (GEFs), including proteins containing Sec7 domains (inhibited by BFA), GBF1 (inhibited by Golgicide A) and BIG1/2 (inhibited by ExoII) (Fig. 6E) ^52^. As expected, treatment of both proliferative and differentiated keratinocytes for 6-24 h with BFA or Golgicide A (GCA) led to the dispersion of Golgi membranes (Figs. 6F & S6E), and inhibition of total protein secretion as shown by SUnSET assay (Fig. 2B), and loss of cell surface binding of WGA lectin (Fig. S6G). Interestingly, BFA and GCA treatment of differentiated keratinocytes concomitantly resulted in the disappearance of typical lysosomes, with LAMP1 labeling only very small punctate structures (0.1 - 0.3 μm) (Fig. 6F) as observed in cells expressing ARF1^T31N^-GFP. Similar results were obtained upon siRNA-mediated depletion of GBF1 (Fig. 6G). In contrast, treatment of differentiated keratinocytes with ExoII led to the dispersion of the Golgi but not to the loss of lysosomal morphology (Fig. 6F), which suggests that the lysosomes are not directly dependent on the late Golgi function. A similar effect was obtained upon BFA, GCA or ExoII treatment of proliferative keratinocytes (Fig. S6E), but not of HeLa cells or primary fibroblasts (not shown). Interestingly, removal of BFA in differentiated keratinocytes, both the lysosomes and the Golgi reappeared with similar kinetics within 3 hours, suggesting a similar sensitivity of the lysosomes to the functional inhibitors of the Golgi (not shown). Together, these data support the role of functional Golgi or intra Golgi function mediated by GBF1-ARF1 in maintaining keratinocyte lysosomes.

## Discussion

In this study, we demonstrate the critical contribution of the Golgi apparatus in the maintenance and functional specialization of lysosomes during keratinocyte differentiation. The specialized properties of keratinocyte lysosomes are denoted by their physical apposition/contact with the Golgi apparatus, the presence of the Golgi tether GRASP65 on the lysosome membrane and the requirement for GRASP65 in maintaining lysosomal morphology and function, the extensive accumulation a *trans*-Golgi enzyme and secretory cargoes in the lysosome lumen, cell surface secretion, and the dependence of lysosome maintenance on Golgi function. We propose that lysosomes in differentiated keratinocytes are specialized for both degradative and secretory functions, and thus are dependent on Golgi association for their function.

The fundamental trafficking mechanisms that regulate unique aspects of intracellular organelle biogenesis and homeostasis in epidermal keratinocytes remain largely unknown. A unique feature of differentiated keratinocytes described here is the organization of the Golgi apparatus as dispersed mini-stacks rather than the characteristic centrally organized ribbon seen in most cell types. Although redistribution of the Golgi apparatus was reported in differentiating epidermal sublayers *in vivo* and differentiating keratinocytes *in vitro* ^37,53^, the molecular mechanism underlying this unique distribution remains unknown. While GRASP55 and GRASP65 are thought to play a key role in organizing the Golgi ribbon morphology ^9,54^, we found that GRASP55 and GRASP65 expression levels were similar in differentiated keratinocytes and proliferative keratinocytes. Moreover, overexpression of either GRASP55 or GRASP65 in differentiated keratinocytes did not reverse the dispersed phenotype. Previous studies in other cellular systems have documented Golgi dispersion in response to high calcium ^55^, to adhesion loss ^56^, during mitosis ^54^ and neuronal differentiation ^15^, each process involving separate molecular pathways. As keratinocyte differentiation is achieved by high calcium incubation, we hypothesize that GRASP55 phosphorylation, perhaps mediated by calcium-responsive Protein Kinase C-alpha, may be responsible for Golgi dispersion ^55^.

The function of GRASPs, especially GRASP65, is likely remodeled during keratinocyte differentiation. Besides its conventional role in maintaining Golgi ribbon morphology, GRASPs have been shown to function as a tether in multiple cellular conditions especially under cellular stress to facilitate unconventional secretion of transmembrane proteins from autophagosome-like structures ^57^. We show here that in differentiated keratinocytes, GRASP65 mediates the close apposition of dispersed Golgi elements with lysosomes and supports cargo transport from the Golgi to lysosomes. Accordingly, GRASP65 depletion resulted in the loss of Golgi-lysosome apposition and a change in lysosome morphology and function. The giant LAMP1-containing acidic compartments that accumulate in GRASP65-depleted cells are deficient in the lysosomal enzyme cathepsin-D, fail to label for lysosomal small GTPase Arl8B, and accumulate the lysosome substrate glucosylceramide, all suggesting the accumulation of dysfunctional lysosomes. Accumulation of secretory cargoes and the *trans*-Golgi enzyme galT to the lysosomal lumen and secretion of VSVG in the cell surface suggest secretory characteristics of LAMP1-positive lysosomes, similar to most lysosome-related organelles ^8,58^. Based on these results, we hypothesize that GRASP65 functions in differentiated keratinocytes as an active tether to both maintain close contacts between Golgi elements and lysosomes and support the capture of cargo-containing vesicles from the Golgi by the lysosomes for ultimate release to the cell surface. The increased size of the LAMP1-positive lysosomes upon GRASP65 inhibition might reflect either the continual accumulation of cargo despite the loss of degradative function, the accumulation of cargo due to a block in secretion, or both. We hypothesize that the unconventional secretory properties of keratinocyte lysosomes likely to contribute to the higher secretory demand of differentiating keratinocytes in the epidermal sublayers ^59,60^. Although GRASP55 has been implicated in unconventional secretion from autophagosomes ^46,61^, its contribution to the secretory properties of keratinocyte lysosomes is unlikely as they do not show significant association with GRASP55, nor GRASP55 depletion had an effect on lysosome morphology. A complete block in total protein secretion in the presence of the Golgi inhibitor GCA, and inhibition of the major fraction of secretion by the lysosomal inhibitor bafilomycin-A1 suggests that the major part of the Golgi mediated secretion is operated through the lysosomes in differentiated keratinocytes.

Their critical dependence on the Golgi apparatus is an additional characteristic that differentiates the lysosomes of differentiated keratinocytes from lysosomes in other cell types and leads us to label them as specialized. Expression of a dominant negative mutant of ARF1 or treatment with inhibitors of the ARF1 GEF GBF1 result in the loss of LAMP1-positive lysosomes in differentiated keratinocytes, but not in cells that harbor conventional lysosomes such as HeLa cells. Moreover, when expressed in keratinocytes, ARF1-GFP labels small dynamic vesicular structures (possibly COPI vesicles) that closely associate and/or fuse with lysosomes by live cell imaging (Fig. 6A). These dynamic ARF1-containing structures near lysosomes are not observed in HeLa cells or other cell types with conventional lysosomes. Together, these observations emphasize the unique modification of both the Golgi apparatus and lysosomes in differentiated keratinocytes. We propose that lysosomes in differentiated keratinocytes mature through constant and direct Golgi input (lysosomal/Golgi cargoes). As ARF1 and GBF1 are required to generate COPI coated vesicles ^16,20,62–64^, we hypothesize that COPI coated vesicles mediate the transport between the Golgi elements and lysosomes in differentiated keratinocytes and thus contribute to the functional maturation/maintenance of the lysosomes. As GRASP65 facilitates Golgi-lysosome apposition and receives Golgi input (Fig. 6G), it would be interesting to test whether functional coordination exists between ARF1 and GRASP65. We hypothesize that the generation of specialized lysosomes is an adaptation to the nutrient deficient environment in the upper skin layers that likely serves the dual purpose of degradation and secretion required to maintain epidermis homeostasis.

## Material and methods

### Chemicals and fluorescence substrates

All the chemicals and reagents used in this study were commercially sourced as mentioned. LysoTracker Red DND-99 (L7528) and Wheat Germ Agglutinin (WGA)-Alexa Fluor 594 (W11262) were obtained from ThermoFisher Scientific (Invitrogen). Dextran-Fluorescein (70K MW, D1822), brefeldin A (B7651), Golgicide A (G0923) and EXOII (E7159) were procured from Sigma-Aldrich. Trypsin and trypsin neutralizer solution were obtained from ThermoFisher Scientific (Invitrogen). Collagen-I was obtained from BD-Biosciences. N-C6:0-NBD-glucosylceramide was from Matreya LLC, USA. Lipofectamine 3000 and oligofectamine were obtained from ThermoFisher Scientific (Invitrogen).

### Antibodies

Following polyclonal and monoclonal primary antibodies were used in 1:100 or, 1:200 dilutions (abbreviations, m=mouse; h=human and r=rat proteins). Abnova: anti-GALT (PAB31346). BD Biosciences: anti-rGM130 (610822) and anti-hp230/golgin-245 (611281). BioRad: anti-hTGN46 (AHP500G). Cell Signaling Technology: anti-LC3A/B (4108). Developmental Studies Hybridoma Bank: anti-hCD63 (H5C6), anti-hLAMP-1 (H4A3) and anti-hLAMP-2 (H4B4). Invitrogen: anti-LAMP1 (14-1079-80). Merck: anti-hCathepsin D (IM16). Proteintech: Arl8b (13049-1). Santa Cruz biotechnology: anti-GAPDH (sc-25778), anti-hGRASP55 (sc-365602), anti-hGRASP65(sc-365434) and anti-hGBF1 (sc-136240). All secondary antibodies were either from Invitrogen or Jackson Immunoresearch and used in 1:500 dilution.

### Primers, siRNAs and plasmids

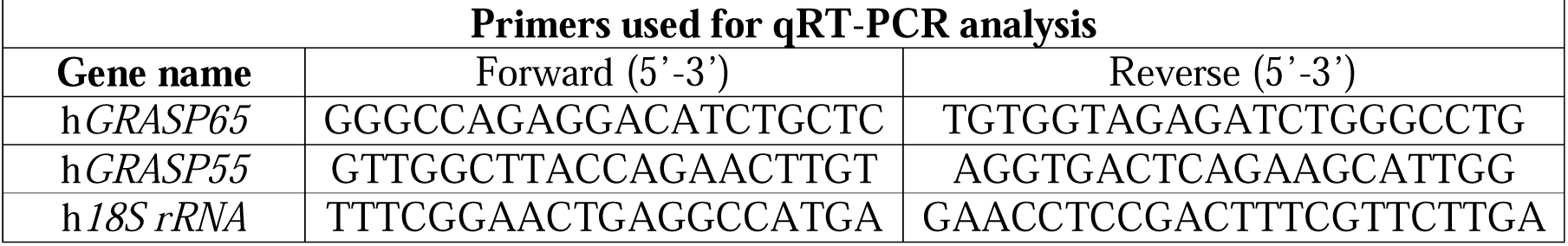

The following target siRNA sequences were synthesized from Eurogentec, Belgium. hGRASP65 (5’-GAUCUCUACCACAGAAUAA-3’), hGRASP55 (5’-GAGUCUGACUGGACUUUCU-3’), hGM130(5’-GAAUCAUGAUGCUGACAAU-3’), hGBF1 (5’- CCUCUGUCAACAAGUUCCU-3’) and control siRNA was purchased from Eurogentec. Following plasmids used in this study were described previously: ARF1^WT^-GFP, ARF1^T31N^-GFP ^65^; GRASP55-GFP, GRASP55-mCherry and GRASP65-GFP ^66^; LAMP1-GFP ^67^; galT-RFP and ManII-GFP ^56^; NPY-Venus ^50^. LAMP1-RFP was obtained from Addgene.

### Primary keratinocyte culture and differentiation

Human neonatal primary keratinocytes were purchased from Lonza, Invitrogen or Life line Cell Technology and grown according to manufacturer’s instructions. Cells were maintained and passaged in the respective medium and used until passage four (P4). For differentiation, cells were grown to 50 - 60% confluence and CaCl_2_ was added to the culture medium to a final concentration of 2 mM. Cells were incubated for 48-60 hours with changing of medium in every 24 h. We have used bigger cell size and dispersal of lysosomes as differentiation characteristics as described previously ^37^.

### Dextran-Fluorescein, LysoTracker Red DND-99 and NBD-Glucosylceramide uptake

Glass coverslips containing control and differentiated keratinocytes were incubated in fresh culture medium with 0.5 mg/ml of dextran-fluorescein for 6 h (included in the total differentiation time) or 50 –75 nM of LysoTracker Red for 10 min in a 37°C incubator with 10% CO_2_. After incubation, the coverslips were washed twice with 1X PBS, fixed with 4% PFA and stained for LAMP1. For NBD-Glucosylceramide (NBD-GCS) uptake, the cells on glass coverslips were first washed with 1XHBSS/HEPES buffer and incubated with a final concentration of 2 μM NBD-GCS in 1XHBSS/HEPES at 4°C for 30 min to promote its binding. After 30 min, the dye was removed, washed twice with HBSS/HEPES buffer and incubated in fresh growth medium for at least 30 min at 37°C in an incubator maintained with 10% CO_2_. Cells were then washed in fresh medium and either imaged in live or, fixed with 4% PFA and imaged.

### Gene expression and siRNA mediated inhibition in primary keratinocytes

For the expression of fluorescent protein tagged constructs, Lipofectamine 3000 was used as per the manufacturer’s instruction. The cells were imaged in between 36 – 48 hours of transfection. Transfection of specific siRNA was performed by using oligofectamine as per the manufacturer’s protocol. Briefly, primary keratinocytes at ∼ 60% confluence were transfected in serum free medium. A final concentration of 20 nM of siRNA was added to the cells and incubated till 72 h. To achieve the differentiation, cells were added with calcium post 24 h of siRNA transfection and then incubated for another 48 h.

### Small molecule treatment and differentiation

Primary keratinocytes were grown on glass coverslips to 50 – 60% confluence. The compounds/drugs or respective solvent (as a control) was added to the medium in duplicates. After 3 hours of treatment, the coverslips were segregated into two sets for control and differentiated conditions. The later set of coverslips was added with CaCl_2_ to a final concentration of 2 mM. All coverslips were incubated for 24h along with the indicated concentrations of compound/solvent to avoid reversal of the phenotype. Cells were then fixed with 4% PFA and continued for immunostaining.

### Gene expression by qRT-PCR analysis

For these experiments, siRNA-mediated gene knockdown was performed in 35 mm dishes. RNA was isolated using GeneJET RNA purification kit (ThermoFisher Scientific) followed by cDNA preparation using cDNA synthesis kit (Fermantas). Same cDNA was used to amplify both the gene of interest and *18S rRNA* in Quant-Studio 6 Flex real-time PCR system (Applied Biosystems). The PCR conditions consisted of AmpliTaq Gold activation at 95°C for 10 min, followed by 40 cycles of denaturation at 95°C for 20s, annealing at 58°C for 25s and extension at 72°C for 30s. A dissociation curve was generated at the end of each cycle to validate the single transcript amplification. The change in SYBR green fluorescence intensity was monitored and then calculated the threshold cycle (C_t_) number. The C_t_ value of the gene was subtracted from respective control to obtain the ΔC_t_ value. The ΔC_t_ value of treated sample was subtracted with the ΔC_t_ value of control to obtain the ΔΔC_t_ value. The gene expression level in the knockdown conditions in compared to the control was expressed as 2^-ΔΔCt^ values from three independent experiments, and plotted as mean±s.e.m. (considering control as 1.00).

### Measuring total protein secretion by SUnSET assay

The assay was performed as described before ^44^ with little modifications. An equal number of primary keratinocytes were grown to ∼ 50% confluency and were either continued as proliferative cells, or differentiated for 48 hours in presence of high calcium. On the day of the experiment cells were cultured in plain medium (without growth factors) with 10μg/ml of puromycin for 30 minutes at 37^0^C. Cells were washed twice with 1XPBS and puromycin, and was chased for 0h and 2h. Cells treated with cycloheximide (CHX;100ng/ml) were added with the drug 30 minutes prior addition of puromycin. While Golgicide A (GCA 10μM final concentration) and or, bafilomycin-A (100nM final concentration) was added along with puromycin. GCA was continued till the whole chasing period. An equal percent of cell lysate or supernatant were loaded and proceeded for immunoblotting. The puromycin signal was detected by using an *anti*-puromycin primary antibody (clone 12D10; Millipore; MABE343). UT= untreated, CHX= cycloheximide, GCA=Golgicide-A, Baf.=Bafilomycin-A1. Total synthesis and secretion are shown by the lysate and in cell medium respectively.

### Immunostaining and immunoblotting

Immunostaining of the cells was performed as described previously ^37^. Briefly, 4% PFA fixed cells were simultaneously permeabilized and immunostained by incubating in the primary antibody solution containing saponin. Cells were incubated with primary antibody at room temperature for approximately 1h followed by incubation with the respective Alexa fluor conjugated secondary antibody for 20-30 min. For WGA-AF594 staining, 4% PFA fixed cells were incubated with 100 μM of WGA-AF594 in 1XPBS for 10 min at RT, followed by staining for LAMP1. For immunoblotting, cell lysate was prepared in SDS lysis buffer and then subjected to immunoblotting analysis as described previously ^37^.

### Fluorescence microscopy

Most of the images including live cell experiments were performed under confocal laser scanning microscope with Airyscan super resolution module (ZEISS LSM 880 with Airyscan) at 63X magnification. Imaging was also done under spinning disk confocal microscope (Inverted Eclipse Ti-E (Nikon), Spinning disk CSU-X1 (Yokogawa)) and, Olympus IX81 motorized inverted fluorescence microscope equipped with a CoolSNAP HQ2 (Photometrics) CCD camera.

### Data analyses in ImageJ/Fiji

Analysis of the cell area, size, numbers and, perimeter of the lysosomes (in μm^2^) was performed by using the watershed function. Lysosome numbers/μm^2^ area was calculated by dividing total number of lysosomes by area of the respective cell. Pearson’s colocalization (*r*), and Mander’s overlap coefficients (M1 and M2) was calculated using JaCoP plugin. Images were captured in Olympus IX81 motorized inverted fluorescence microscope to calculate localization coefficients. The nm distance between the Golgi membrane to lysosome membrane and, Golgi stacks to lysosomes (Figs. 2D and 4C) were measured after respective scale calibration.

### Structured Illumination Microscopy, 3D rendering and, analysis of length and breadth

Structured Illumination Microscopy was performed using ZEISS Elyra 7 with Lattice SIM microscope with a lateral (x-y) resolution of ∼120 nm and an axial (z) resolution of ∼300 nm. Samples were imaged as 3D stack with an axial interval of 200 nm between focal planes and, at 63X magnification. The images were reconstructed and rendered with the help of 4-D viewer in Metamorph 7.0 (Molecular Devices, USA) software.

For length and breadth analysis, Z-stack SIM micrographs of labelled lysosomes (LAMP1-RFP) and Golgi (GRASP65-GFP) were converted to maximum intensity projection. Using average + standard deviation values of fluorescence intensity of entire image, the signal was segmented to a binary image. The segmented image was used with integrated morphometric analysis tool to log length, breadth and area for the objects in Metamorph 7.0 (Molecular Devices, USA). Golgi and lysosomes of 3 differentiated keratinocytes from 3 different experiments were included. The number of organelles analyzed is mentioned on the respective graph. Independent values were plotted as a function of fractions and significance was analyzed by Mann-Whitney non-parametric analysis.

### Volumetric analysis of lysosome and Golgi

Micrographs in Z-stack (imaged in Airyscan super resolution) of labelled lysosomes (LAMP1) and Golgi (GM130), were processed using 4D viewer module in Metamorph 7.0 (Molecular Devices, USA) (Previously reported in ^68,69^). Micrographs were thresolded for intensity to segment the regions for the labelled lysosomes and Golgi for each focal image. With 4D viewer the structure was rendered and reconstructed to generate a 3D model of the Z-stack. Using filtering for size, appropriate lysosome and Golgi models were selected for volumetric analysis. Volume of individual objects detected was logged to CSV files and cumulative distribution was calculated using GraphPad Prism software (version 6.04). The value of significance was analyzed by Mann-Whitney non-parametric analysis between respective groups. Five cells of each category (control siRNA and GRASP65 siRNA treated cells) from 3 independent experiments were included in the analysis.

### Cryo-sectioning, immunogold labelling and Transmission Electron Microscopy

Ultrathin cryo sections were prepared and immunogold labelling was performed as described in ^70^. Briefly, the control and differentiated keratinocytes were fixed with 2% PFA and 0.25% Glutaraldehyde in 0.1 M phosphate buffer solution (pH 7.4) overnight at 4°C. Cells were embedded in 10% gelatin and cryoprotected by using 2.1M sucrose and stored in liquid nitrogen. Ultrathin cryosections (100nm) were prepared with an ultra-cryomicrotome UC7 FCS (Leica). Unlabelled cryosections were either processed for imaging after contrasting with uranyl acetate (as shown in Fig. 2B) or, processed for single/ double immunogold labeling with protein-A conjugated gold particles of 10 nm and 15nm diameter (Cell Microscopy Center, Department of Cell Biology, Utrecht University).Images were acquired with a Transmission Electron Microscope (Tecnai Spirit G2; ThermoFischer Scientific, Eindhoven, The Netherlands) equipped with a 4k CCD camera (Orius 1000, Gatan Inc., Pleasanton, CA). For all analyses ≥ 150 images from at least two different experiments were included.

### Statistical analysis in graph pad prism (version 6.04)

Cells from at least three independent experiments were included for statistical analysis (number of cells ‘n’ is mentioned on the respective graph). Pearson’s colocalization coefficient ‘*r*’, and Mander’s overlap coefficients M1 and M2 (mentioned together as ‘localization coefficient’) values were plotted as mean±s.e.m. and the statistical significance was calculated by unpaired two tailed Student’s *t*-test. The non-normal distribution of Golgi to lysosome distances (Figs. 2D and 4C was analyzed by frequency distribution analysis. The value of significance was calculated either by Mann Whitney non-parametric test (between two samples) or by Kolmogorov-Smirnov non parametric test to analyze cumulative distributions specified in the respective sections. \**P*≤0.05, \*\**P* ≤ 0.01, \*\*\**P* ≤ 0.001, \*\*\*\**P* ≤ 0.0001 and ns = non-significant. Graphs in ‘red’ color indicates control and ‘blue’ indicates differentiated conditions.

## Author contributions

S.M., S.R.G.S., G.R. and B.G. designed research. S.M. performed all the experiments (unless specified), analyzed data, coordinated between co-authors and, wrote the manuscript. P.B. performed electron microscopy experiments. V.B., L.E. and D.N. performed super resolution microscopy and data analysis. N.G. performed a part of immunofluorescence data quantification and provided technical help. G.R., D.N. and B.G. gave critical inputs during manuscript preparation. S.R.G.S and G.R. edited the manuscript. All the authors read and approved the final version of the manuscript.

## Acknowledgement

We are grateful to Satyajit Mayor (NCBS), Yanzhuang Wang (University of Michigan), Franck Perez (Inst. Curie, Paris, France), Nagaraj Balasubramanian (IISER Pune, India) and Amitabha Majumdar (HUL, Bangalore, India) for generous gifts of constructs and reagents. We thank Sarit Agasti, JNCASR for allowing us access Structured Illumination Microscope. We sincerely thank Cedric Delevoye (Institute Curie, Paris) for insightful discussions and critical reading of the manuscript. Kavya Gopal is acknowledged for technical help during the study.

We sincerely acknowledge IISc divisional imaging facility. We acknowledge the Nikon Imaging Center @ Institut Curie-CNRS and the PICT-IBiSA. EM sample preparation and cryosectioning at structure and membrane compartments, UMR144, Institute Curie is greatly acknowledged. We acknowledge Adrien Candat and the IBENS electron imaging facility (IMACHEM-IBiSA), member of the French National Research infrastructure France-BioImaging.

## Funding

This work is supported by the DBT-Wellcome Trust India Alliance ECF (IA/E/17/503685) to S.M. SRGS acknowledges Science and Engineering Research Board (CRG/2019/000281), Department of Biotechnology (BT/PR4982/AGR/36/718/2012 and BT/PR32489/BRB/10/1786/2019), DBT-NBACD (BT/HRD-NBA-NWB/38/2019-20).

## Conflict of interest

The authors declare that they have no conflict of interest.

## Supplementary Figures

**Figure S1. Comparative analyses of the lysosomes of proliferative and differentiated keratinocytes.** (**A**) Time-lapse imaging of differentiated keratinocyte expressing LAMP1-RFP by Airyscan super-resolution microscope. Arrowheads following the dynamics of small LAMP1 carriers. Two different events from the same cell are presented (top and bottom panel). Scale bar=2.5 μm. (**B**-**C**) IFM analysis of proliferative and differentiated keratinocytes co-immunostained for LAMP1 with either LAMP2 (B) or LysoTracker Red (C). Cells were visualized under Airyscan super-resolution microscope. Insets represent 2.5 times magnified white boxed regions. Arrowheads point to the colocalization between the markers. White stars represent the LAMP1 structures that are negative for LysoTracker Red. Scale bar=10 μm.

**Figure S2. Presence of Golgi lysosome apposition in differentiated keratinocytes.** (**A**) Loading controls for the SUnSET assay shown in Fig. 2B. The total synthesis and secretion were measured with or without the presence of small molecules. P= proliferative and D = differentiated conditions. UT= untreated, CHX= cycloheximide, GCA= Golgicide-A. Total synthesis and secretion is shown by the lysate and in cell medium/supernatant respectively. (**B-D**) Proliferative and differentiated keratinocytes were labeled for (B)WGA-alexa flour 594 (C) Mitotracker red or, (D) EEA1, and co-labelled for LAMP1. Cells were visualized by Airyscan super-resolution microscope at 60X magnification. Insets in (B) represent 2.5 times magnified white boxed region. Scale bar=10 μm. **(E)** IFM analysis of proliferative and differentiated keratinocytes co-stained for GM130 and p230 (first panel); or LAMP1 with either GM130, p230, TGN46 or expressed ARF1-GFP. Cells were visualized by Airyscan super-resolution microscope. The numbers (1-3) on the differentiated keratinocytes represent the magnified regions that are presented in Fig. 2C. Insets represent 2.5 times magnified white boxed regions. Pearson’s correlation coefficient (*r*) and Mander’s overlap coefficients (M1 and M2) in proliferative cells were calculated and plotted as mean±s.e.m. **** on the graphs indicates significant difference between M1 and M2 values. n=number of cells used in the quantification method. Scale bar=10 μm. (**F**) LAMP1-immunogold labelling of ultrathin cryosections to define lysosomes. Represented lysosomal structures are present in both, proliferative and differentiated keratinocytes. Scale bar=200 nm. (**G**) Comparative Golgi morphology in proliferative and differentiated keratinocytes. Scale bar=200 nm. Please note that the LAMP1 immunogold labelled sections were also used to analyze the Golgi morphology. Negative staining of the Golgi apparatus by LAMP1 confirms the specificity of LAMP1 immunolabelling in 2F. (**F**) proliferative and differentiated keratinocytes were co-immuno stained for LAMP1 with either LAMP2 (red=proliferative; blue=differentiated) or DAPI respectively. The localization coefficients (Pearson’s correlation coefficient (*r*) and Mander’s overlap coefficients, (M1 and M2) values were plotted as mean±s.e.m. n=the number of cells used for quantifications.

**Figure S3. GRASP65 surrounds the lysosomes in differentiated keratinocytes.** (**A**) Differentiated keratinocytes expressing GRASP65-GFP and GRASP55-mCherry were imaged by a spinning disk microscope. Arrows and arrowheads represent respectively to the colocalization of GRASP65 with GRASP55 in dissociated Golgi stacks and, unique ring like structures of GRASP65 that are negative for GRASP55. Insets represent 2.5 times magnified white boxed regions. Scale bar=10μm. (**B**) mRNA expressions of GRASP55 and GRASP65 in proliferative and differentiated keratinocytes was measured by semi quantitative-RT PCR. (**C**) Live cell imaging of proliferative keratinocyte expressing GRASP65-GFP and LAMP1-RFP by Airyscan super-resolution microscope. Scale bar=10 μm. (**D**) The graph represents % of proliferative cells having at least one GRASP65-GFP ring in live or in steady state. n=number of cells. (**E**-**F**) Quantification of coefficients (Pearson’s correlation coefficient, *r* and Mander’s overlap coefficients, M1 and M2) between GRASP65-GFP and LAMP1-RFP in the proliferative keratinocytes in steady (E) and, in live (F) conditions. The values were plotted as mean±s.e.m. n=number of cells. (**G**) IFM analysis of proliferative and differentiated keratinocytes stained for LAMP1 and LC3 (G) arrowheads represent to the LC3-positive autophagosomes. (**H**) Co-immunostaining of endogenous GRASP65 with LAMP1. Cells were imaged by Airyscan super-resolution microscope. Insets represent 2.5 times magnified white boxed regions. Scale bar=10 μm.

**Figure S4. GRASP65 mediates Golgi-lysosome contacts in differentiated keratinocytes. (A)** IFM analysis of the control/GRASP65 siRNA treated proliferative keratinocytes using GRASP65 and LAMP1 co-immunostaining. (**B-D**) IFM analysis of the respective siRNA treated differentiated keratinocytes (as labelled). Cells were co immunostained for LAMP1 with either GRASP65 (B), or GRASP55 (C), or GM130 (D) and imaged by Airyscan super resolution microscope. Scale bar=10 μm. Insets represent 2.5 times magnified white boxed regions. Scale bar=5 μm. (**E**) Transcript analysis of GRASP65 and 55 in respective siRNA treated condition, plotted as fold change. Values of 3 independent experiments were included. (**F**) Immunoblotting analysis of control siRNA and GRASP65 siRNA treated cells. Blots were probed with GRAPS65 or beta-actin (as internal control). (**G)** Characterization of the giant LAMP1-positive structures in GRASP65siRNA treated differentiated keratinocytes. Cells were stained for LAMP1 and Cathepsin-D, or Arl8B (1^st^ and 2^nd^ panels). In the 3^rd^ panel, cells were incubated with NBD-glucosylceramide (GC) and lysotracker red. Scale bar=10 μm. The merged insets are presented in Fig. 4D. **(H)** Volumetric distribution analysis was performed as described in materials and methods. The log10 values were plotted as cumulative fractions. Statistical significance of the volume of the Golgi in control siRNA and GRASP65 siRNA treated cells were calculated by Mann-Whitney non parametric test.

**Figure S5. Differentiated keratinocytes produce specialized secretory lysosomes** (**A**) IFM analysis of differentiated keratinocytes expressing ManII-GFP and LAMP1-RFP. Cells were imaged by Airyscan super-resolution microscope. Insets from two different regions (white boxes) were magnified into 2.5 times and shown separately. Scale bar=10 μm. (**B**) IFM analysis of the co-expressed ts045VSVG-GFP and LAMP1-RFP. Transfected cells were incubated at 40^0^C overnight followed by synchronized release at 32^0^C over the time points of 30 min, 60 min and 90 min. Cells were imaged by a Spinning disk microscope at 60X magnification. Scale bar=10 μm. (**C**) Total protein secretion from the proliferative and differentiated keratinocytes was measured by the SUnSET assay in presence or absence of the lysosomal inhibitor bafilomycin-A1. The total secretion was measured at 0h and, 2h. P= proliferative and D = differentiated conditions. UT= untreated, GCA= Golgicide-A., Baf. =bafilomycin-A1. Total synthesis and secretion are shown by the lysate and in cell medium/supernatant respectively. (**D**) IFM analysis of differentiated keratinocytes expressing LAMP1-GFP and galT-RFP in presence of bafilomycin-A1(50nM). Imaging was done by Airyscan microscope at 60X magnification. Scale bar=10 μm.

**Figure S6. A functional Golgi maintains keratinocyte lysosomes.** (**A**) Time lapse analysis of proliferative keratinocytes expressing ARF1^WT^-GFP and LAMP1-RFP. Live cell imaging was performed by Airyscan super-resolution microscope. Scale bar=10μm. (**B**) localization coefficients (Pearson’s correlation coefficient, *r* and Mander’s overlap coefficients, M1 and M2) in proliferative cells between ARF1-GFP and LAMP1-RFP were plotted as mean±s.e.m. (**C**) IFM analysis of Golgi morphology of differentiated keratinocytes expressing ARF1^T31N^-GFP, immuno stained either for GM130 or GRASP65. Scale bar=10 μm. (**D**) analysis of lysosome morphology of proliferative keratinocyte co-expressed with ARF1^T31N^-GFP and LAMP1-RFP. Scale bar=10 μm. (**E**) Proliferative keratinocytes were treated with pharmacological inhibitors as described in materials and methods. Cells were fixed and stained for GM130 and LAMP1. Insets represent 2.5 times magnified white boxed regions. Arrows indicate to the Golgi and lysosome morphology in all treatment conditions. Scale bar=10 μm. (**F**) HeLa cells were transfected with either ARF1-GFP or ARF1^T31N^-GFP and immunostained with LAMP1. Scale bar=10 μm. (**G**) Proliferative and differentiated keratinocytes either treated with DMSO or BFA, fixed and stained for WGA Alexa flour 594 (WGA-A594) and LAMP1. Insets represent 2.5 times magnified white boxed regions. Arrows indicate to the effect of BFA on the plasma membrane binding of WGA. In all images: scale bar=10 μm.

## Supplementary Videos

**Video S1:** Live cell imaging of differentiated keratinocyte expressing LAMP1-RFP. The scale bar and time frames are mentioned in the video.

**Video S2:** Live cell imaging of differentiated keratinocytes expressing GRASP65-GFP and LAMP1-RFP. The scale bar and time frames are mentioned in the video.

**Video S3:** Live cell imaging of proliferative keratinocyte expressing GRASP65-GFP and LAMP1-RFP. The scale bar and time frames are mentioned in the video.

**Video S4:** Live cell imaging of differentiated keratinocyte expressing ARF1-GFP and LAMP1-RFP. The scale bar and time frames are mentioned in the video.

**Video S5:** Live cell imaging of proliferative keratinocyte expressing ARF1-GFP and LAMP1-RFP expressed. The scale bar and time frames are mentioned in the video.

## Notes

### Competing Interest Statement

The authors have declared no competing interest.

### Summary of Updates

We have significantly updated the result section. However, the scientific conclusion remains the same.

## References

1. Bravo-Sagua, R., Torrealba, N., Paredes, F., Morales, P.E., Pennanen, C., Lopez-Crisosto, C., Troncoso, R., Criollo, A., Chiong, M., Hill, J.A., et al. (2014). Organelle communication: signaling crossroads between homeostasis and disease. Int J Biochem Cell Biol 50, 55–59. 10.1016/j.biocel.2014.01.019.

2. Atakpa, P., Thillaiappan, N.B., Mataragka, S., Prole, D.L., and Taylor, C.W. (2018). IP3 Receptors Preferentially Associate with ER-Lysosome Contact Sites and Selectively Deliver Ca(2+) to Lysosomes. Cell Rep 25, 3180–3193 e3187. 10.1016/j.celrep.2018.11.064.

3. Wong, Y.C., Kim, S., Peng, W., and Krainc, D. (2019). Regulation and Function of Mitochondria-Lysosome Membrane Contact Sites in Cellular Homeostasis. Trends Cell Biol 29, 500–513. 10.1016/j.tcb.2019.02.004.

4. Chu, B.B., Liao, Y.C., Qi, W., Xie, C., Du, X., Wang, J., Yang, H., Miao, H.H., Li, B.L., and Song, B.L. (2015). Cholesterol transport through lysosome-peroxisome membrane contacts. Cell 161, 291–306. 10.1016/j.cell.2015.02.019.

5. Frohlich, F., and Gonzalez Montoro, A. (2023). The role of lysosomes in lipid homeostasis. Biol Chem 404, 455–465. 10.1515/hsz-2022-0287.

6. Luzio, J.P., Poupon, V., Lindsay, M.R., Mullock, B.M., Piper, R.C., and Pryor, P.R. (2003). Membrane dynamics and the biogenesis of lysosomes. Mol Membr Biol 20, 141–154. 10.1080/0968768031000089546.

7. Wartosch, L., Bright, N.A., and Luzio, J.P. (2015). Lysosomes. Curr Biol 25, R315–316. 10.1016/j.cub.2015.02.027.

8. Delevoye, C., Marks, M.S., and Raposo, G. (2019). Lysosome-related organelles as functional adaptations of the endolysosomal system. Curr Opin Cell Biol 59, 147–158. 10.1016/j.ceb.2019.05.003.

9. Xiang, Y., and Wang, Y. (2010). GRASP55 and GRASP65 play complementary and essential roles in Golgi cisternal stacking. J Cell Biol 188, 237–251. 10.1083/jcb.200907132.

10. Xiang, Y., Zhang, X., Nix, D.B., Katoh, T., Aoki, K., Tiemeyer, M., and Wang, Y. (2013). Regulation of protein glycosylation and sorting by the Golgi matrix proteins GRASP55/65. Nat Commun 4, 1659. 10.1038/ncomms2669.

11. Ayala, I., Mascanzoni, F., and Colanzi, A. (2020). The Golgi ribbon: mechanisms of maintenance and disassembly during the cell cycle. Biochem Soc Trans 48, 245–256. 10.1042/BST20190646.

12. Rambourg, A., Clermont, Y., Chretien, M., and Olivier, L. (1993). Modulation of the Golgi apparatus in stimulated and nonstimulated prolactin cells of female rats. Anat Rec 235, 353–362. 10.1002/ar.1092350304.

13. Krause, W.J. (2000). Brunner’s glands: a structural, histochemical and pathological profile. Prog Histochem Cytochem 35, 259–367.

14. Thayer, D.A., Jan, Y.N., and Jan, L.Y. (2013). Increased neuronal activity fragments the Golgi complex. Proc Natl Acad Sci U S A 110, 1482–1487. 10.1073/pnas.1220978110.

15. Wei, J.H., and Seemann, J. (2017). Golgi ribbon disassembly during mitosis, differentiation and disease progression. Curr Opin Cell Biol 47, 43–51. 10.1016/j.ceb.2017.03.008.

16. Donaldson, J.G., Honda, A., and Weigert, R. (2005). Multiple activities for Arf1 at the Golgi complex. Biochimica et Biophysica Acta (BBA) - Molecular Cell Research 1744, 364–373. 10.1016/j.bbamcr.2005.03.001.

17. Satoh, A., Hasegawa, Y., and Honjo, Y. (2015). The Roles of GRASP55/65 in Golgi Formation and Function. Trends in Glycoscience and Glycotechnology 27, 33–36. 10.4052/tigg.27.33.

18. Wang, Y., Satoh, A., and Warren, G. (2005). Mapping the functional domains of the Golgi stacking factor GRASP65. J Biol Chem 280, 4921–4928. 10.1074/jbc.M412407200.

19. Zhang, X., and Wang, Y. (2015). GRASPs in Golgi Structure and Function. Front Cell Dev Biol 3, 84. 10.3389/fcell.2015.00084.

20. Spang, A. (2002). ARF1 regulatory factors and COPI vesicle formation. Curr Opin Cell Biol 14, 423–427. 10.1016/s0955-0674(02)00346-0.

21. Wang, Y., Wei, J.H., Bisel, B., Tang, D., and Seemann, J. (2008). Golgi cisternal unstacking stimulates COPI vesicle budding and protein transport. PLoS One 3, e1647. 10.1371/journal.pone.0001647.

22. Scott, K.L., Kabbarah, O., Liang, M.C., Ivanova, E., Anagnostou, V., Wu, J., Dhakal, S., Wu, M., Chen, S., Feinberg, T., et al. (2009). GOLPH3 modulates mTOR signalling and rapamycin sensitivity in cancer. Nature 459, 1085–1090. 10.1038/nature08109.

23. Thomas, J.D., Zhang, Y.J., Wei, Y.H., Cho, J.H., Morris, L.E., Wang, H.Y., and Zheng, X.F. (2014). Rab1A is an mTORC1 activator and a colorectal oncogene. Cancer Cell 26, 754–769. 10.1016/j.ccell.2014.09.008.

24. Makhoul, C., and Gleeson, P.A. (2021). Regulation of mTORC1 activity by the Golgi apparatus. Fac Rev 10, 50. 10.12703/r/10-50.

25. Wang, T., and Hong, W. (2002). Interorganellar regulation of lysosome positioning by the Golgi apparatus through Rab34 interaction with Rab-interacting lysosomal protein. Mol Biol Cell 13, 4317–4332. 10.1091/mbc.e02-05-0280.

26. Starling, G.P., Yip, Y.Y., Sanger, A., Morton, P.E., Eden, E.R., and Dodding, M.P. (2016). Folliculin directs the formation of a Rab34-RILP complex to control the nutrient-dependent dynamic distribution of lysosomes. EMBO Rep 17, 823–841. 10.15252/embr.201541382.

27. Zhang, X., and Wang, Y. (2018). GRASP55 facilitates autophagosome maturation under glucose deprivation. Mol Cell Oncol 5, e1494948. 10.1080/23723556.2018.1494948.

28. Gotz, T.W.B., Puchkov, D., Lysiuk, V., Lutzkendorf, J., Nikonenko, A.G., Quentin, C., Lehmann, M., Sigrist, S.J., and Petzoldt, A.G. (2021). Rab2 regulates presynaptic precursor vesicle biogenesis at the trans-Golgi. J Cell Biol 220. 10.1083/jcb.202006040.

29. Lie, P.P.Y., Yang, D.S., Stavrides, P., Goulbourne, C.N., Zheng, P., Mohan, P.S., Cataldo, A.M., and Nixon, R.A. (2021). Post-Golgi carriers, not lysosomes, confer lysosomal properties to pre-degradative organelles in normal and dystrophic axons. Cell Rep 35, 109034. 10.1016/j.celrep.2021.109034.

30. Hennings, H., Kruszewski, F.H., Yuspa, S.H., and Tucker, R.W. (1989). Intracellular calcium alterations in response to increased external calcium in normal and neoplastic keratinocytes. Carcinogenesis 10, 777–780. 10.1093/carcin/10.4.777.

31. Bikle, D.D., Xie, Z., and Tu, C.L. (2012). Calcium regulation of keratinocyte differentiation. Expert Rev Endocrinol Metab 7, 461–472. 10.1586/eem.12.34.

32. Akinduro, O., Sully, K., Patel, A., Robinson, D.J., Chikh, A., McPhail, G., Braun, K.M., Philpott, M.P., Harwood, C.A., Byrne, C., et al. (2016). Constitutive Autophagy and Nucleophagy during Epidermal Differentiation. J Invest Dermatol 136, 1460–1470. 10.1016/j.jid.2016.03.016.

33. Mahanty, S., and Setty, S.R.G. (2021). Epidermal Lamellar Body Biogenesis: Insight Into the Roles of Golgi and Lysosomes. Front Cell Dev Biol 9, 701950. 10.3389/fcell.2021.701950.

34. Yoshihara, N., Ueno, T., Takagi, A., Oliva Trejo, J.A., Haruna, K., Suga, Y., Komatsu, M., Tanaka, K., and Ikeda, S. (2015). The significant role of autophagy in the granular layer in normal skin differentiation and hair growth. Arch Dermatol Res 307, 159–169. 10.1007/s00403-014-1508-0.

35. Monteleon, C.L., Agnihotri, T., Dahal, A., Liu, M., Rebecca, V.W., Beatty, G.L., Amaravadi, R.K., and Ridky, T.W. (2018). Lysosomes Support the Degradation, Signaling, and Mitochondrial Metabolism Necessary for Human Epidermal Differentiation. J Invest Dermatol 138, 1945–1954. 10.1016/j.jid.2018.02.035.

36. Bochenska, K., Moskot, M., Malinowska, M., Jakobkiewicz-Banecka, J., Szczerkowska-Dobosz, A., Purzycka-Bohdan, D., Plenkowska, J., Slominski, B., and Gabig-Ciminska, M. (2019). Lysosome Alterations in the Human Epithelial Cell Line HaCaT and Skin Specimens: Relevance to Psoriasis. Int J Mol Sci 20. 10.3390/ijms20092255.

37. Mahanty, S., Dakappa, S.S., Shariff, R., Patel, S., Swamy, M.M., Majumdar, A., and Setty, S.R.G. (2019). Keratinocyte differentiation promotes ER stress-dependent lysosome biogenesis. Cell Death Dis 10, 269. 10.1038/s41419-019-1478-4.

38. Pols, M.S., van Meel, E., Oorschot, V., ten Brink, C., Fukuda, M., Swetha, M.G., Mayor, S., and Klumperman, J. (2013). hVps41 and VAMP7 function in direct TGN to late endosome transport of lysosomal membrane proteins. Nat Commun 4, 1361. 10.1038/ncomms2360.

39. Bright, N.A., Davis, L.J., and Luzio, J.P. (2016). Endolysosomes Are the Principal Intracellular Sites of Acid Hydrolase Activity. Curr Biol 26, 2233–2245. 10.1016/j.cub.2016.06.046.

40. Johnson, D.E., Ostrowski, P., Jaumouille, V., and Grinstein, S. (2016). The position of lysosomes within the cell determines their luminal pH. J Cell Biol 212, 677–692. 10.1083/jcb.201507112.

41. Barral, D.C., Staiano, L., Guimas Almeida, C., Cutler, D.F., Eden, E.R., Futter, C.E., Galione, A., Marques, A.R.A., Medina, D.L., Napolitano, G., et al. (2022). Current methods to analyze lysosome morphology, positioning, motility and function. Traffic 23, 238–269. 10.1111/tra.12839.

42. Bekier, M.E., 2nd, Wang, L., Li, J., Huang, H., Tang, D., Zhang, X., and Wang, Y. (2017). Knockout of the Golgi stacking proteins GRASP55 and GRASP65 impairs Golgi structure and function. Mol Biol Cell 28, 2833-2842. 10.1091/mbc.E17-02-0112.

43. Goodman, C.A., and Hornberger, T.A. (2013). Measuring protein synthesis with SUnSET: a valid alternative to traditional techniques? Exerc Sport Sci Rev 41, 107–115. 10.1097/JES.0b013e3182798a95.

44. Fourriere, L., Kasri, A., Gareil, N., Bardin, S., Bousquet, H., Pereira, D., Perez, F., Goud, B., Boncompain, G., and Miserey-Lenkei, S. (2019). RAB6 and microtubules restrict protein secretion to focal adhesions. J Cell Biol 218, 2215–2231. 10.1083/jcb.201805002.

45. Klumperman, J., and Raposo, G. (2014). The complex ultrastructure of the endolysosomal system. Cold Spring Harb Perspect Biol 6, a016857. 10.1101/cshperspect.a016857.

46. Giuliani, F., Grieve, A., and Rabouille, C. (2011). Unconventional secretion: a stress on GRASP. Curr Opin Cell Biol 23, 498–504. 10.1016/j.ceb.2011.04.005.

47. Zhang, X., and Wang, Y. (2020). Nonredundant Roles of GRASP55 and GRASP65 in the Golgi Apparatus and Beyond. Trends Biochem Sci 45, 1065–1079. 10.1016/j.tibs.2020.08.001.

48. Schaub, B.E., Berger, B., Berger, E.G., and Rohrer, J. (2006). Transition of galactosyltransferase 1 from trans-Golgi cisterna to the trans-Golgi network is signal mediated. Mol Biol Cell 17, 5153–5162. 10.1091/mbc.e06-08-0665.

49. Grigoriev, I., Splinter, D., Keijzer, N., Wulf, P.S., Demmers, J., Ohtsuka, T., Modesti, M., Maly, I.V., Grosveld, F., Hoogenraad, C.C., and Akhmanova, A. (2007). Rab6 regulates transport and targeting of exocytotic carriers. Dev Cell 13, 305–314. 10.1016/j.devcel.2007.06.010.

50. Patwardhan, A., Bardin, S., Miserey-Lenkei, S., Larue, L., Goud, B., Raposo, G., and Delevoye, C. (2017). Routing of the RAB6 secretory pathway towards the lysosome related organelle of melanocytes. Nat Commun 8, 15835. 10.1038/ncomms15835.

51. Lachuer, H., Le, L., Leveque-Fort, S., Goud, B., and Schauer, K. (2023). Spatial organization of lysosomal exocytosis relies on membrane tension gradients. Proc Natl Acad Sci U S A 120, e2207425120. 10.1073/pnas.2207425120.

52. Manolea, F., Claude, A., Chun, J., Rosas, J., and Melancon, P. (2008). Distinct functions for Arf guanine nucleotide exchange factors at the Golgi complex: GBF1 and BIGs are required for assembly and maintenance of the Golgi stack and trans-Golgi network, respectively. Mol Biol Cell 19, 523–535. 10.1091/mbc.e07-04-0394.

53. Yamanishi, H., Soma, T., Kishimoto, J., Hibino, T., and Ishida-Yamamoto, A. (2019). Marked Changes in Lamellar Granule and Trans-Golgi Network Structure Occur during Epidermal Keratinocyte Differentiation. J Invest Dermatol 139, 352–359. 10.1016/j.jid.2018.07.043.

54. Colanzi, A., and Corda, D. (2007). Mitosis controls the Golgi and the Golgi controls mitosis. Curr Opin Cell Biol 19, 386–393. 10.1016/j.ceb.2007.06.002.

55. Ireland, S., Ramnarayanan, S., Fu, M., Zhang, X., Zhang, J., Li, J., Emebo, D., and Wang, Y. (2020). Cytosolic Ca(2+) Modulates Golgi Structure Through PKCalpha-Mediated GRASP55 Phosphorylation. iScience 23, 100952. 10.1016/j.isci.2020.100952.

56. Singh, V., Erady, C., and Balasubramanian, N. (2018). Cell-matrix adhesion controls Golgi organization and function through Arf1 activation in anchorage-dependent cells. J Cell Sci 131. 10.1242/jcs.215855.

57. Rabouille, C., and Linstedt, A.D. (2016). GRASP: A Multitasking Tether. Front Cell Dev Biol 4, 1. 10.3389/fcell.2016.00001.

58. Bowman, S.L., Bi-Karchin, J., Le, L., and Marks, M.S. (2019). The road to lysosome-related organelles: Insights from Hermansky-Pudlak syndrome and other rare diseases. Traffic 20, 404–435. 10.1111/tra.12646.

59. Boyce, S.T. (1994). Epidermis as a secretory tissue. J Invest Dermatol 102, 8–10. 10.1111/1523-1747.ep12371721.

60. Katz, A.B., and Taichman, L.B. (1994). Epidermis as a secretory tissue: an in vitro tissue model to study keratinocyte secretion. J Invest Dermatol 102, 55–60. 10.1111/1523-1747.ep12371732.

61. Nickel, W. (2010). Pathways of unconventional protein secretion. Curr Opin Biotechnol 21, 621–626. 10.1016/j.copbio.2010.06.004.

62. Kawamoto, K., Yoshida, Y., Tamaki, H., Torii, S., Shinotsuka, C., Yamashina, S., and Nakayama, K. (2002). GBF1, a guanine nucleotide exchange factor for ADP-ribosylation factors, is localized to the cis-Golgi and involved in membrane association of the COPI coat. Traffic 3, 483–495. 10.1034/j.1600-0854.2002.30705.x.

63. Nebenfuhr, A., Ritzenthaler, C., and Robinson, D.G. (2002). Brefeldin A: deciphering an enigmatic inhibitor of secretion. Plant Physiol 130, 1102–1108. 10.1104/pp.011569.

64. Yang, J.S., Valente, C., Polishchuk, R.S., Turacchio, G., Layre, E., Moody, D.B., Leslie, C.C., Gelb, M.H., Brown, W.J., Corda, D., et al. (2011). COPI acts in both vesicular and tubular transport. Nat Cell Biol 13, 996–1003. 10.1038/ncb2273.

65. Kumari, S., and Mayor, S. (2008). ARF1 is directly involved in dynamin-independent endocytosis. Nat Cell Biol 10, 30–41. 10.1038/ncb1666.

66. Zhang, X., Wang, L., Lak, B., Li, J., Jokitalo, E., and Wang, Y. (2018). GRASP55 Senses Glucose Deprivation through O-GlcNAcylation to Promote Autophagosome-Lysosome Fusion. Dev Cell 45, 245–261 e246. 10.1016/j.devcel.2018.03.023.

67. Marwaha, R., Arya, S.B., Jagga, D., Kaur, H., Tuli, A., and Sharma, M. (2017). The Rab7 effector PLEKHM1 binds Arl8b to promote cargo traffic to lysosomes. J Cell Biol 216, 1051–1070. 10.1083/jcb.201607085.

68. Venkatachalapathy, M., Belapurkar, V., Jose, M., Gautier, A., and Nair, D. (2019). Live cell super resolution imaging by radial fluctuations using fluorogen binding tags. Nanoscale 11, 3626–3632. 10.1039/c8nr07809b.

69. Kedia, S., Ramakrishna, P., Netrakanti, P.R., Singh, N., Sisodia, S.S., Jose, M., Kumar, S., Mahadevan, A., Ramanan, N., Nadkarni, S., and Nair, D. (2021). Alteration in synaptic nanoscale organization dictates amyloidogenic processing in Alzheimer’s disease. iScience 24, 101924. 10.1016/j.isci.2020.101924.

70. Hurbain, I., Romao, M., Bergam, P., Heiligenstein, X., and Raposo, G. (2017). Analyzing Lysosome-Related Organelles by Electron Microscopy. Methods Mol Biol 1594, 43–71. 10.1007/978-1-4939-6934-0_4.

