## Supplementary figures and images for "Biogenesis of specialized lysosomes in differentiated keratinocytes relies on close apposition with the Golgi apparatus"

### Supplementary Figures

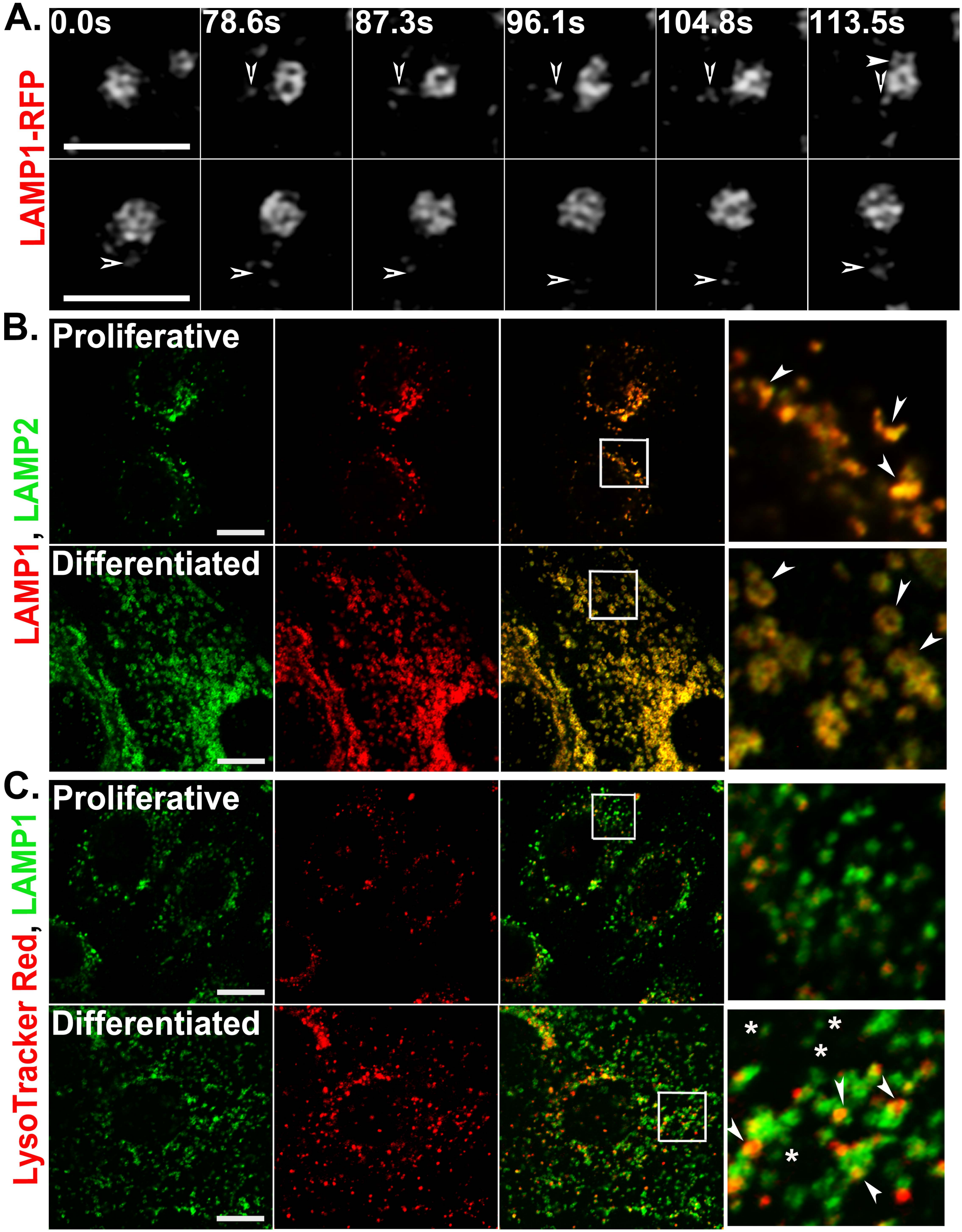

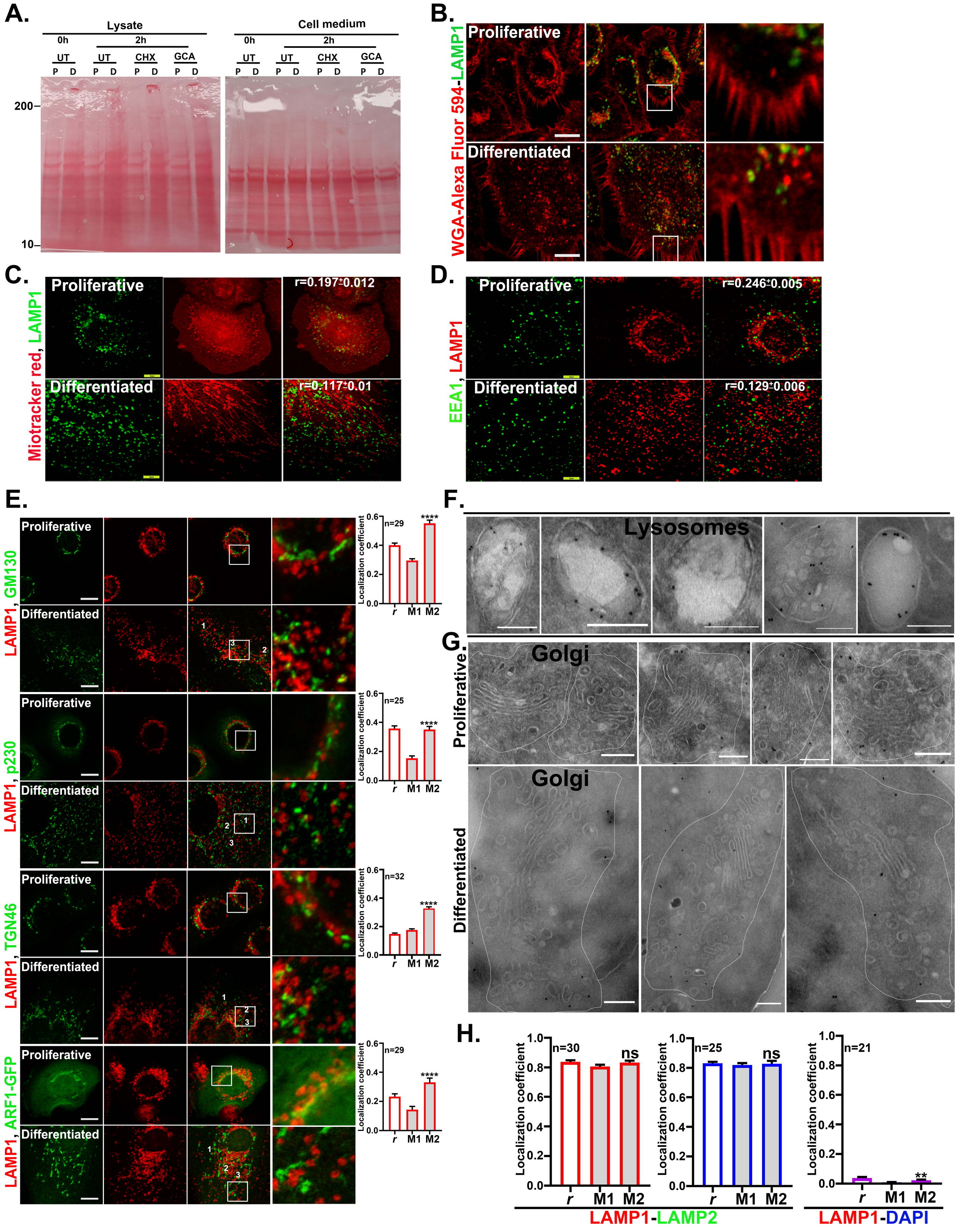

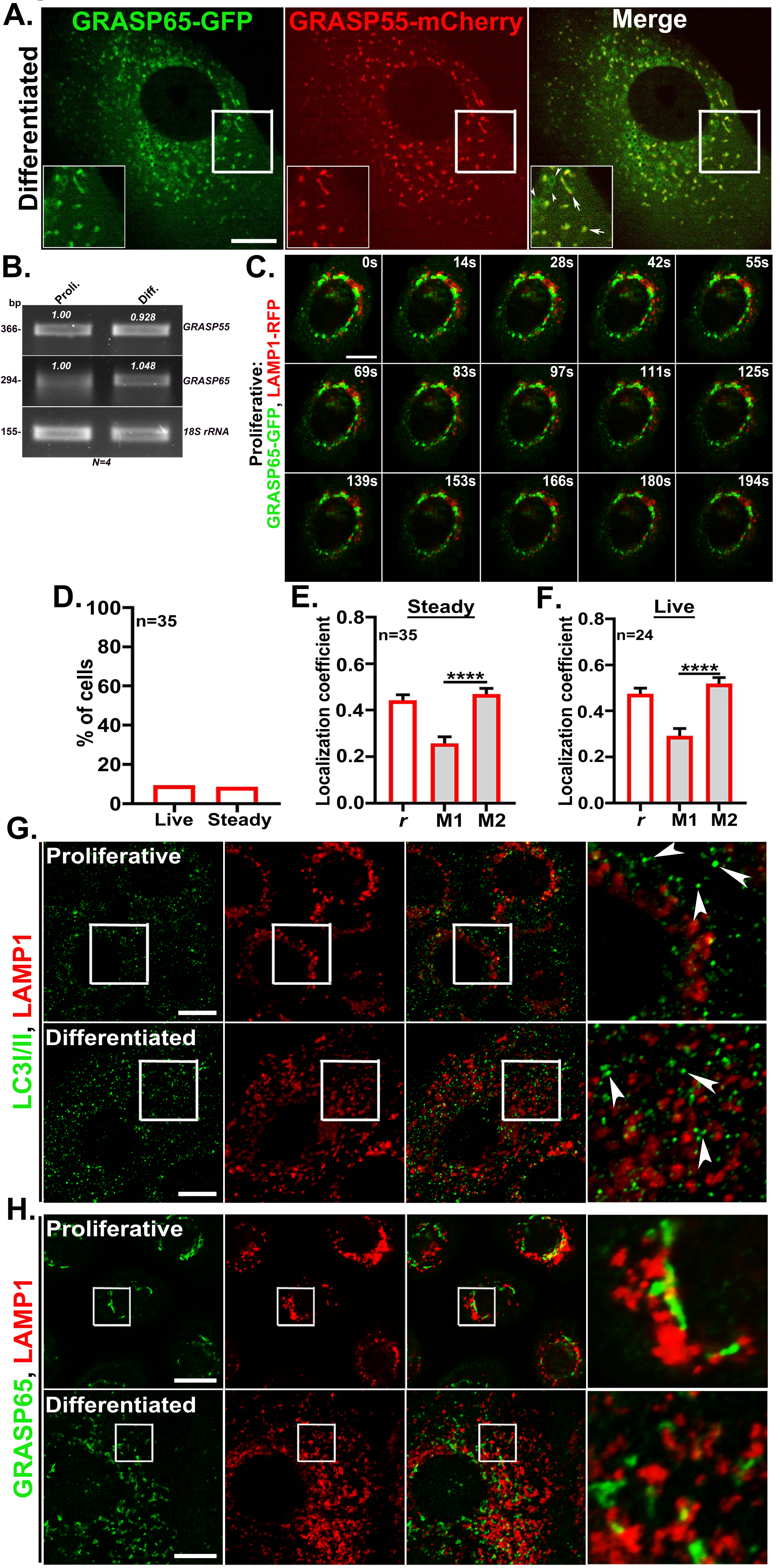

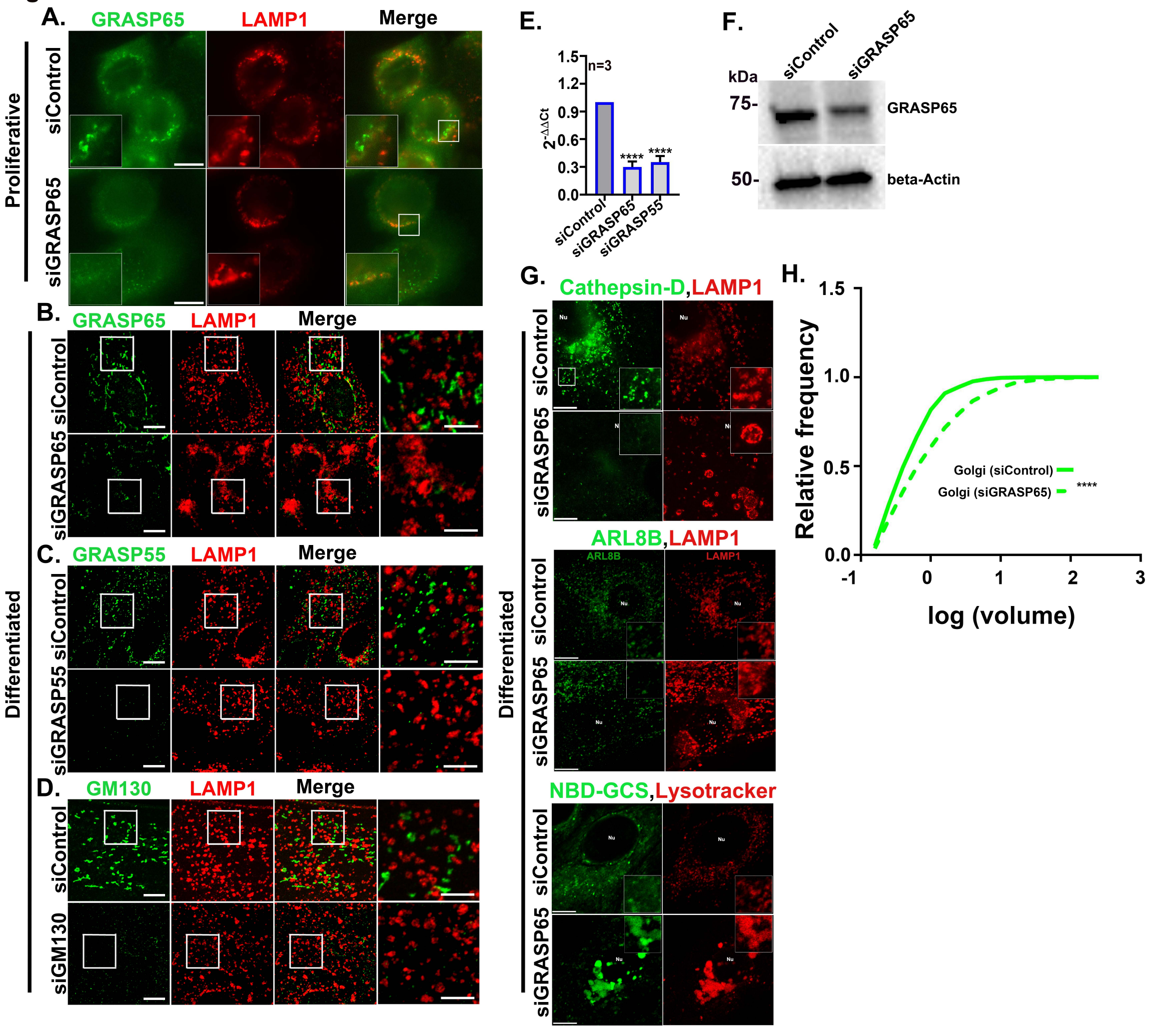

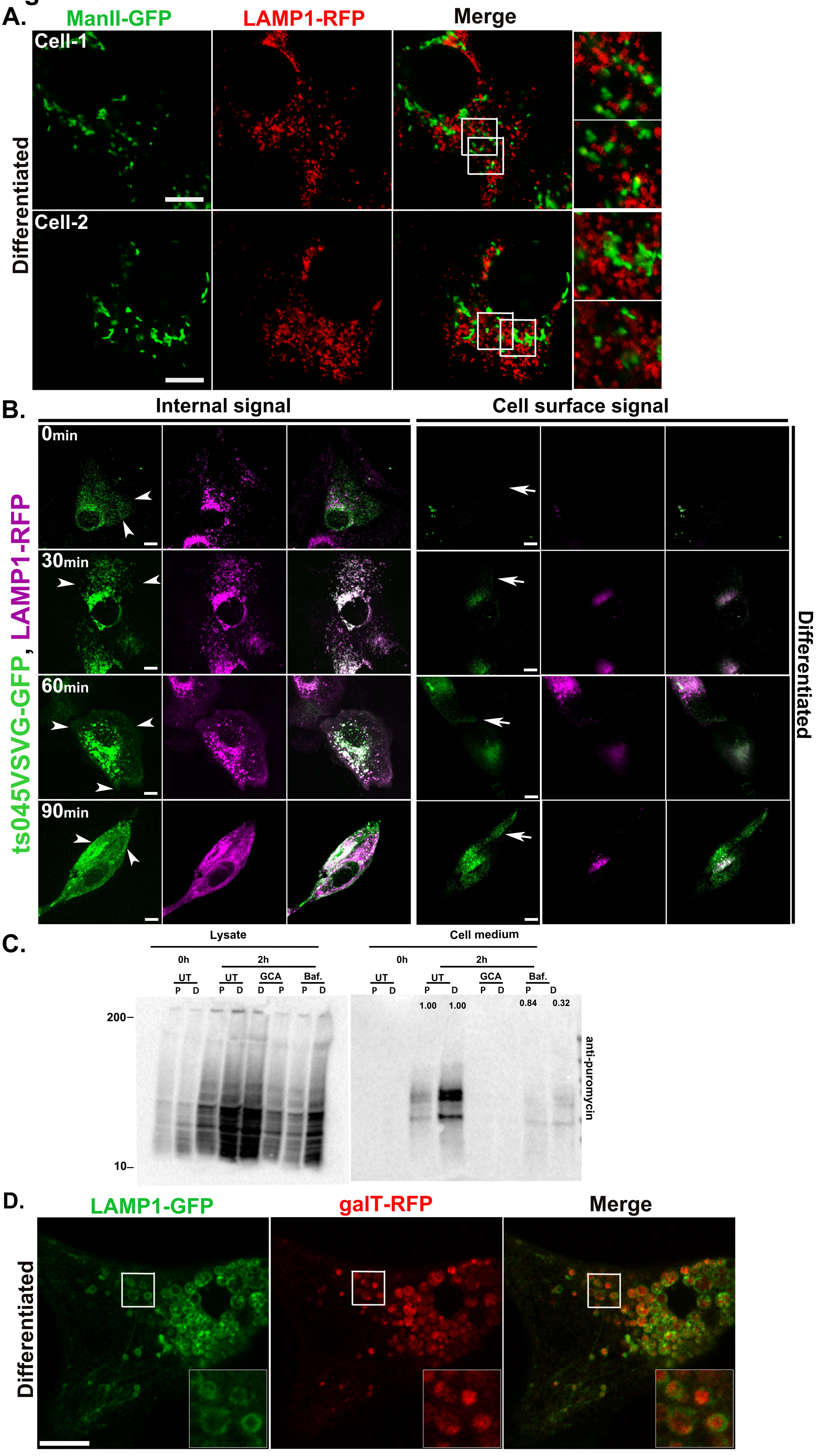

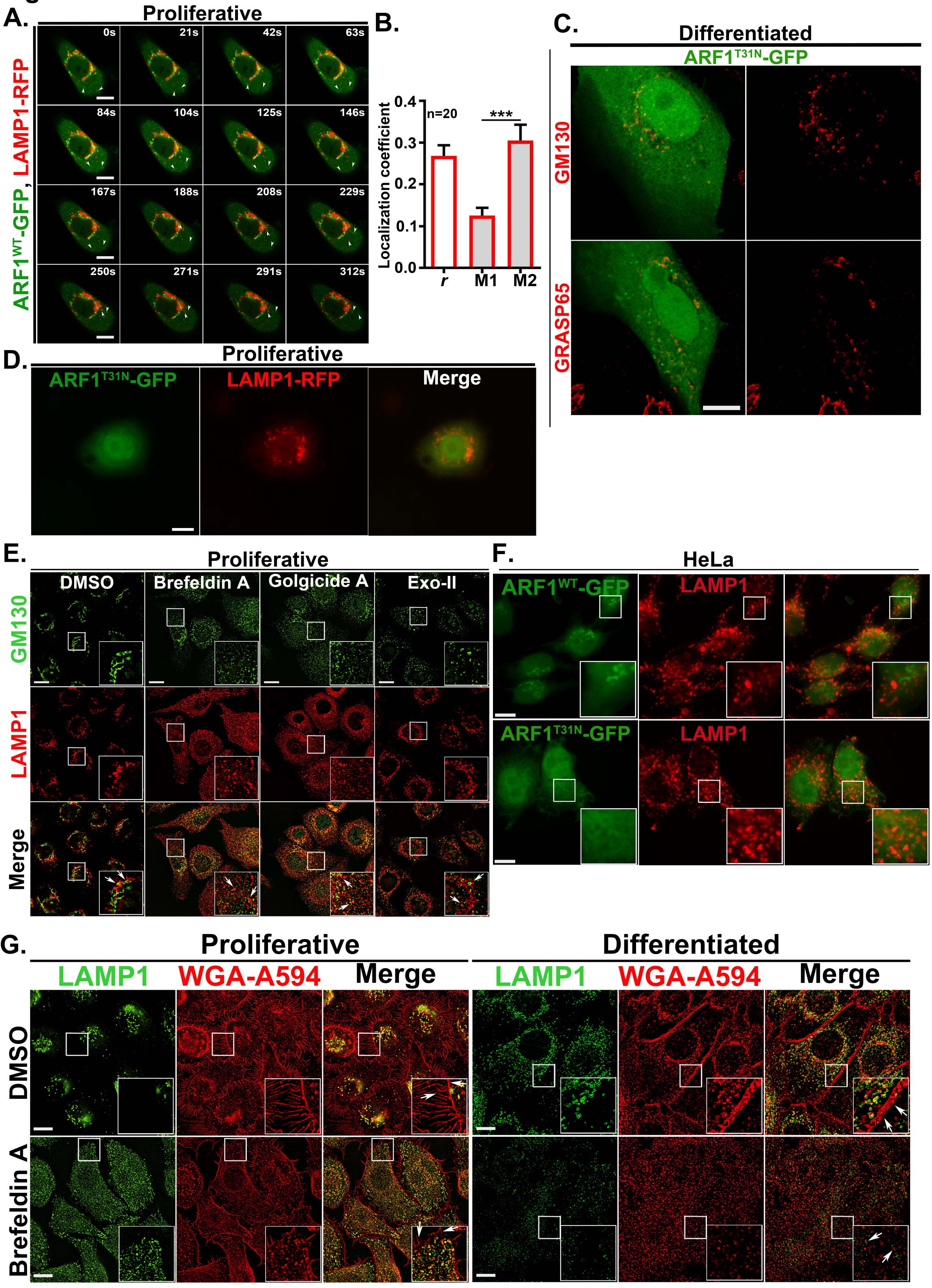
